# Motor usage imprints microtubule stability in the shaft

**DOI:** 10.1101/2021.04.09.439170

**Authors:** Mireia Andreu-Carbó, Simon Fernandes, Marie-Claire Velluz, Karsten Kruse, Charlotte Aumeier

## Abstract

Tubulin dimers assemble into dynamic microtubules which are used by molecular motors as tracks for intracellular transport. Organization and dynamics of the microtubule network is commonly thought to be regulated at the polymer ends, where tubulin-dimers can be added or removed. Here we show that molecular motors running on microtubules cause exchange of dimers along the shaft. These sites of dimer exchange act as rescue sites where depolymerising microtubules stop shrinking and start re-growing. Consequently, the average length of microtubules increases depending on how frequently they are used as motor tracks. An increase of motor activity densifies the cellular microtubule network and enhances cell polarity. Running motors leave marks in the shaft serving as traces of microtubule usage to organize the polarity landscape of the cell.

## INTRODUCTION

Kinesins transport cargo (Verhey et al., 2011) and can also affect microtubule dynamics (Drummond, 2011). Microtubules are 25 nm wide tubes, polymerising at the tip by addition of α−β GTP-tubulin heterodimers. After incorporation at the plus-tip, GTP-tubulin dimers are hydrolysed (Mitchison & Kirschner, 1984). If GDP-tubulin dominates at the tip, then the microtubule experiences a catastrophe, they switch from a growing to a shrinking phase. While shrinking, tubulin dimers are removed from the plus-end. Recent reports showed that not only the tip, but also the rest of the microtubule, its ‘shaft’, shows dynamic behaviour (Aumeier et al., 2016; de Forges et al., 2016; Dimitrov et al., 2008; Reid et al., 2017; Schaedel et al., 2015, 2019; Vemu et al., 2018). Tubulin can dissociate from the shaft spontaneously (Schaedel et al., 2019), or upon mechanical stress (Schaedel et al., 2015), resulting in damage sites along the GDP-shaft. These damage sites get repaired by incorporating GTP-tubulin. Some proteins have recently been reported to cause tubulin dissociation from the shaft or to mediate GTP-tubulin incorporation (Vemu et al., 2018; Triclin et al., 2018; Aher et al., 2020;). At a GTP-tubulin incorporation site, depolymerizing microtubules can be rescued and start re-growing. Therefore, sites of microtubule damage and dimer exchange are constructive areas leading to an increased microtubule lifetime (Aumeier et al., 2016).

By acting at the plus-end, kinesins can change the catastrophe rate (Arellano-Santoyo et al., 2017; Walczak et al., 1996) as well as the polymerisation (Hibbel et al., 2015; J Howard & Hyman, 2003, 2007) and depolymerisation speeds (Arellano-Santoyo et al., 2017; J Howard & Hyman, 2003, 2007; Hunter et al., 2003), but no effect on the rescue rate has been previously reported *in vitro*. Binding of kinesin-1 to the microtubule shaft can change the microtubule conformation. Microtubules crowded with at least 10 % kinesin-1 show increased axial spacing between GDP-tubulin subunits, which resembles the elongated structure of a GTP-microtubule without changing the nucleotide state (Muto et al., 2005; Peet et al., 2018; Shima et al., 2018). Once bound to a microtubule, a kinesin-1 molecule runs across hundreds of tubulin dimers without dissociating (Block et al., 1990; Hancock & Howard, 1998). In a cell, kinesin-1 transports various cargos by binding preferentially to the subpopulation of microtubules that are acetylated and/or detyrosinated (Cai et al., 2009). These post-translational modifications have been proposed as underlying mechanisms for preferential binding (Cai et al., 2009; Hammond et al., 2010), but *in vitro* experiments showed only slight effects on the affinity of kinesin-1 for tubulin with posttranslational modifications (Kaul et al., 2014; Soppina et al., 2012).

While the role of microtubule tip-regulation is well established (Mitchison & Kirschner, 1984), regulation of the shaft in cells is largely unexplored. We previously showed that a local increase in shaft dynamics ultimately causes directed migration towards the site of the cell where shaft dynamics is increased and thereby the microtubule lifetime is prolonged (Aumeier et al., 2016).

Here we report that kinesin-1 activity regulates the lifetime and length of its microtubule-track by increasing shaft dynamics. We recently reported in a non-dynamic assay that the minus-end directed motor Klp2 causes tubulin dissociation from end-stabilised microtubules, which destabilises mechanically microtubules in vitro unless tubulin incorporation is allowed (Triclin et al., 2018). Beyond an effect on the mechanic stability of microtubules in a static system, we show here by using dynamic *in vitro* assays and cells that kinesin-1 running on microtubules control microtubule network dynamics through rescue in the shaft. The increase in shaft dynamics is causing more rescue events, which result in a closer proximity between microtubule tips and the plasma membrane. We show that the effect of kinesin-1 on microtubule shaft dynamics ultimately results in a network asymmetry increasing the peripheral microtubule network.

## RESULTS

### Kinesin-1 induces tubulin incorporation along the microtubule shaft in a concentration dependent manner

To test whether running kinesins could induce tubulin-incorporation along the microtubule in a concentration dependent manner, we purified kinesin-1-GFP and performed an assay to monitor microtubule repair (Figure S1A). Using our previously reported two-colour assay (Aumeier et al., 2016), we polymerised microtubules from green Atto488-tubulin dimers and stabilized the tip; then, for 15 minutes, we added free red Atto565-tubulin dimers which can continue to polymerise at the tip or get incorporated along the shaft. This allows us to monitor the incorporation sites, which appear along the shaft as red spots embedded in otherwise green microtubules (Figure 1A, and for important details and controls of this assay see Figure S1A).

**Figure 1.**
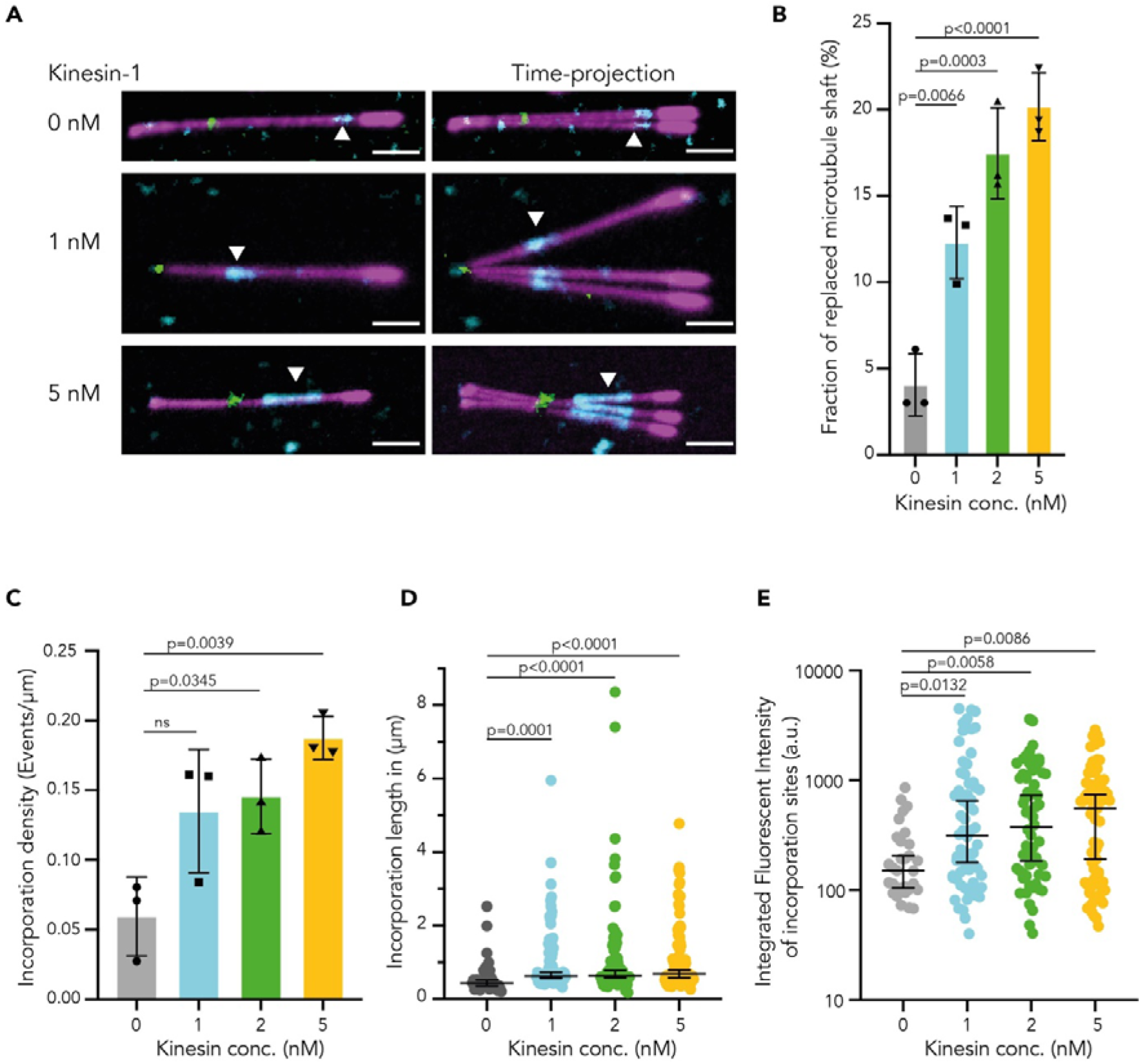
Kinesin-1 induces tubulin incorporation along the microtubule shaft. **(A)** Tubulin incorporation sites (cyan) along GDP-microtubules (magenta; 20% labelled) grown from a seed (green) and capped by GMPCPP (magenta; 70% labelled) in presence of 0 nM, 1 nM, and 5 nM kinesin-1. White arrowheads: incorporation sites. Time projection (max intensity) of 4 subsequent images of single microtubules. Scale bar 2 µm. **(B)** Fraction of replaced microtubule shaft (total length of all incorporation sites divided by the total microtubule length). Mean fraction in % with SD. Statistics: one-way ANOVA. **(C)** Density of incorporation sites (total number of incorporation sites divided by total microtubule length). Mean density with SD. Statistics: one-way ANOVA. For each condition in B and C, three independent experiments were analysed with a total of N= 128, total analysed length L: 821 µm (Control); N= 138, L: 1260 µm (1 nM kinesin-1); N= 135, L: 620 µm (2 nM); N= 139, L: 1055 µm (5nM) microtubules. **(D)** Length of incorporation sites. Median with 95% CI. For each condition, three independent experiments were analysed with a total of N= 45 (Control), 155 (1 nM kinesin-1), 143 (2 nM), and 201 (5 nM) incorporation sites. Statistics: Kruskal-Wallis test. **(E)** Integrated fluorescence intensity of incorporation sites. Median intensity values with 95% CI. Statistics: Kruskal-Wallis test. For each condition three independent experiments were analysed with a total of N= 31 (Control), 64 (1 nM kinesin-1), 60 (2 nM), and 69 (5 nM) incorporation sites.

While running on microtubules (Figures S1B and S1C), kinesin-1 affected GTP-tubulin incorporation into the shaft. In the absence of kinesin-1, spontaneous tubulin incorporation sites covered only 4 % of the microtubules after 15 minutes (Figure 1B). Upon 15 minutes incubation with 5 nM running kinesin-1, microtubule renewal was strongly enhanced: tubulin incorporation sites now covered 20% of the microtubules (Figure 1B). This increase is mainly due to a three-fold increase of the density of incorporation sites along the shaft (Figure 1C). Whereas the density of incorporation sites increased in the presence of running kinesin-1 in a concentration dependent manner (Figure 1C), the length of individual sites increased only slightly with the motor concentration (Figure 1D). However, within incorporation sites the number of exchanged dimers increased with increasing kinesin-1 concentration (Figure 1E). Taken together these results show that kinesin-1 caused defects in the microtubule shaft which were repaired by incorporation of GTP-tubulin dimers at a frequency that was considerably higher than the spontaneous frequency.

### Running Kinesin-1 increase microtubule rescue frequency

Incorporation sites at the shaft have the potential to act as rescue sites (Aumeier et al., 2016; de Forges et al., 2016; Vemu et al., 2018). To address whether the kinesin-1-mediated incorporation sites affect microtubule rescue, we looked at microtubule dynamics in the presence of running kinesin-1 (Figures S2A and S2B). We grew microtubules from GMPCPP-stabilized microtubule seeds in presence of 8 µM tubulin and 41 mM KCl, which yielded dynamic microtubules with an average length of 3.4 ±2.3 µm (n=344 microtubules, Figure 2A). Typically, shrinking microtubules disassembled completely and only in one fourth of the cases (85/384) a rescue event was observed (Figure 2A). After addition of kinesin-1, the shrinking velocity and the catastrophe frequency were not affected, and also the growth velocity remained comparable to the control (Figures 2B-E, S2C and S2B). In contrast, the rescue frequency increased up to four-fold with increasing kinesin-1 concentration (Figure 2D). In the presence of 2 nM kinesin-1, more than half of the shrinking microtubules were rescued (188/309). Therefore, the increased rescue events due to running kinesin-1 correlated with an increase in incorporation site frequency, consistent with previous observations showing that incorporation sites are rescue sites (Aumeier et al., 2016; de Forges et al., 2016; Vemu et al., 2018).

**Figure 2.**
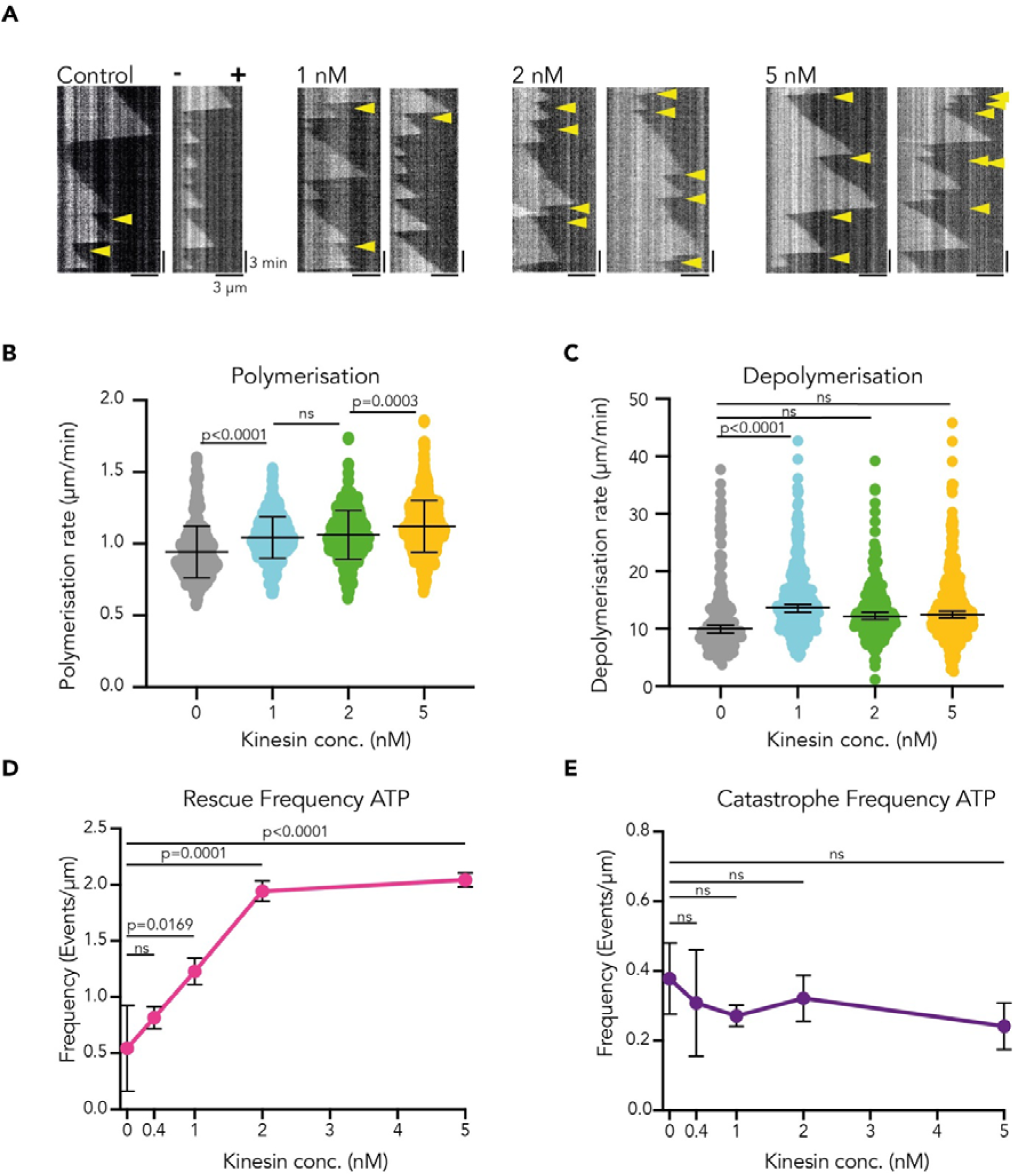
Running kinesin-1 increases microtubule rescue frequency. **(A)** Representative kymographs of microtubule dynamics. Microtubules were polymerised from seeds in the presence of 8 µM tubulin and different kinesin-1 concentrations (0 nM, 1 nM, 2 nM, and 5 nM). Yellow arrowheads indicate rescue events. Plus-ends are oriented to the right. **(B)** Polymerisation speed of microtubules in presence of different kinesin-1 concentrations. Statistics: one-way ANOVA. Mean with SD. **(C)** Depolymerisation speed of microtubules at different kinesin-1 concentrations. Statistics: two-way ANOVA with nested group in log transformed data. Median with 95% CI. In B and C, the polymerisation and depolymerisation speeds were measured for three independent experiments with 296 (Control), 473 (1 nM kinesin-1), 307 (2 nM), and 528 (5 nM) microtubules in total. **(D)** Rescue frequency as a function of kinesin-1 concentration. **(E)** Catastrophe frequency as a function of kinesin-1 concentration. In D and E rescue and catastrophe frequency were measured for each condition in 3-4 independent experiments with 389 (Control), 214 (0.4 nM kinesin-1), 661 (1 nM), 321 (2 nM), and 557 (5 nM) microtubules in total. Statistics: one-way ANOVA. Mean with SD.

### Increase in microtubule rescue only happens if kinesin-1 runs

In the presence of the non-hydrolysable ATP analogue AMP-PNP, kinesins bind to microtubules, but do not run (Hancock & Howard, 1999). In these conditions, the unbinding rate was also reduced and microtubules became densely covered by motors (Figure S3A). Immotile kinesin-1 covering microtubules did not change the microtubule catastrophe frequency or depolymerisation behaviour (Figures S3B, S3C and Video S1). In contrast to our assay with motile kinesin-1, the rescue frequency was not increased in the presence of immotile kinesin-1, indicating that kinesin-1 binding by itself cannot generate rescue sites. In fact, the dense coverage of microtubules by immotile motors led to a decrease in the rescue frequency and in the incorporation frequency (Figures S3B and S3D). This indicates that immotile kinesin-1 could protect the shaft from spontaneous tubulin exchange. Therefore, only running kinesin-1 cause rescue sites.

### Single Kinesin-1 runs can cause rescue events

To study the effect of single kinesin runs on the microtubule rescue frequency, we used 0.4 nM kinesin-1, a concentration which generates isolated, non-colliding runs on microtubules (Figure S2E). Under these conditions, the microtubule rescue frequency was already increased 1.5-fold with respect to controls without kinesin-1 (Figure 2D). At higher kinesin-1 concentrations, the rescue frequency increased linearly with the motor concentration showing that the effect of more kinesins was merely additive. Cooperative effects between kinesin-1 molecules were therefore unlikely to affect the rate of rescue. We conclude that tubulin incorporation along the microtubule can be triggered by single kinesin-1 molecules.

### Kinesin-1 causes tubulin exchange rarely

How often does a single molecule of kinesin-1 lead to an incorporation site? After 15 minutes, the density of incorporation sites in the presence of 2 nM kinesin-1 was 0.14 ± 0.027 events/µm of which 0.06 ± 0.028 events/µm correspond to spontaneous events. Therefore, 2 nM kinesin-1 caused 0.08 incorporation-events per µm. Considering the number of motors binding to microtubules (5.2 kinesin-1/µm within 15 min), and the average kinesin-1 run length (1.4 ± 0.5 µm, Figure S1C), an incorporation site occurred every 91 µm along a kinesin-1-track. This implies that an incorporation site was induced every 11300 kinesin-1 steps. We conclude that kinesin-1 causes incorporation sites very rarely, but that these rare events can change considerably the rescue frequency (Figure 2D).

To study if the rare tubulin incorporation events induced by running kinesins are in principle sufficient to explain our observed increased rescue frequencies, we performed stochastic simulations of the system. In the simulations, a microtubule is effectively described as a polymer represented by a dynamic linear lattice (Johann et al., 2012; Melbinger et al., 2012), with sites that are either in a ‘GTP’- or ‘GDP’-state (Video S2, Figure S4A and see Methods). In sites where there is GTP, a rescue event can occur. We obtained the values for our simulation parameters from our *in vitro* experiments (Table S1). Only the probability at which a shrinking polymer rescues at a GTP-site could not be measured experimental and was therefore used as a fit parameter. The rescue frequency emerging from the simulations matched that observed experimentally in the absence of motor particles (Figures 3A and 3B).

**Figure 3.**
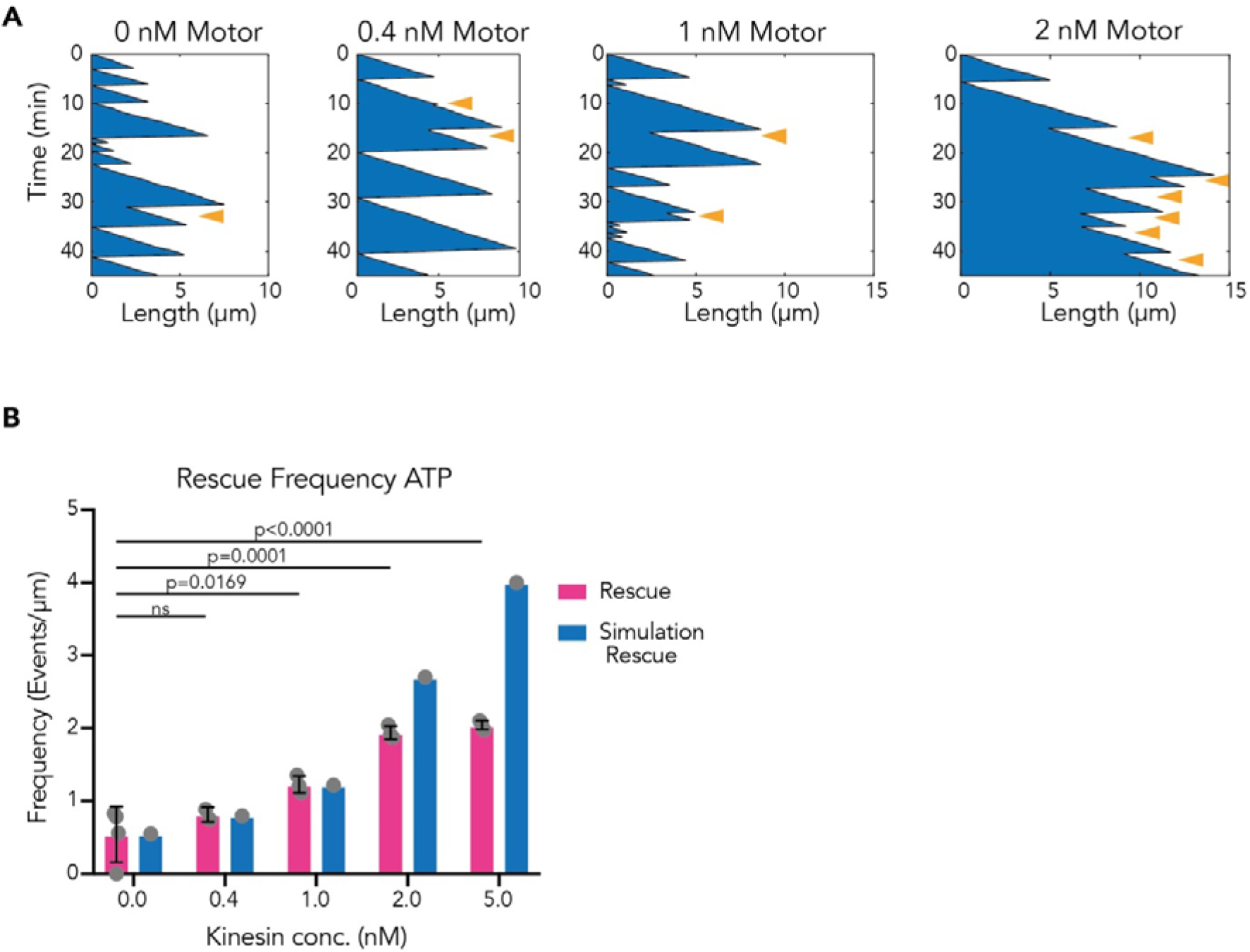
Numerical simulation: Running motors increase rescue frequency. **(A)** Representative kymographs of polymer dynamics in presence of different motor particle attachment rates corresponding to different kinesin-1 concentrations in the experiments (0 nM, 0.4 nM, 1 nM, 2 nM). Orange arrowheads indicate rescue events. **(B)** Comparison of experimental rescue frequency data as in Figure 2D (pink bars) with polymer rescue frequency at steady state obtained from simulations (blue bars). Note that the experiments for 2 and 5 nM kinesin-1 had not yet reached steady state.

To clarify the impact of kinesin-1 on tubulin incorporation along the shaft, we introduced motor particles in our simulation (Johann et al., 2012; Melbinger et al., 2012). Our data showed that motor particles moving on the lattice can change the state of individual lattice sites from GDP to GTP with a given probability p_X_ (Video S3, Figure S4B and see Methods). For p_X_=1.8×10^−9^, we reproduced in our simulations the kinesin-1-dependent rescue frequency observed experimentally. In our simulations, increasing the number of motor particles running on the polymer increased the rescue frequency in a concentration dependent manner, as we have observed experimentally (Figures 3A and 3B). Therefore, rare incorporations caused by running motors are still frequent enough to account for a considerable increase in rescue events as seen in our experiments.

### Running kinesin-1 causes unlimited growth of microtubules

We noted in our simulations an increase in the average polymer length when motor particles were present (Figure 3A). The length increased with the number of motor particles moving on the lattice (Figures 4A and S4C). The density of motor particles on the polymer can be adjusted by changing their binding rate in the simulation. We found that depending on the number of motor particles, the average polymer growth can be bound (i.e. reach a plateau) or unlimited (Figure 4A). In the simulations, at low motor particle density, the rescue frequency was low and growth was bound, because, following a catastrophe, many polymers depolymerised completely (70%). Thus, polymerisation and depolymerisation were balanced and a steady-state with constant average polymer length was reached. In contrast, at high motor particle density, which caused a high rescue frequency, complete depolymerisation could be observed only at initial stages of the simulation. In this scenario, frequent rescue events extended the polymer by adding length to a pre-existing lattice, leading to unlimited growth (Dogterom & Leibler, 1993). This unlimited growth regime occurred above a threshold value of motor particles which corresponded to the density observed experimentally at 2 nM kinesin-1.

**Figure 4.**
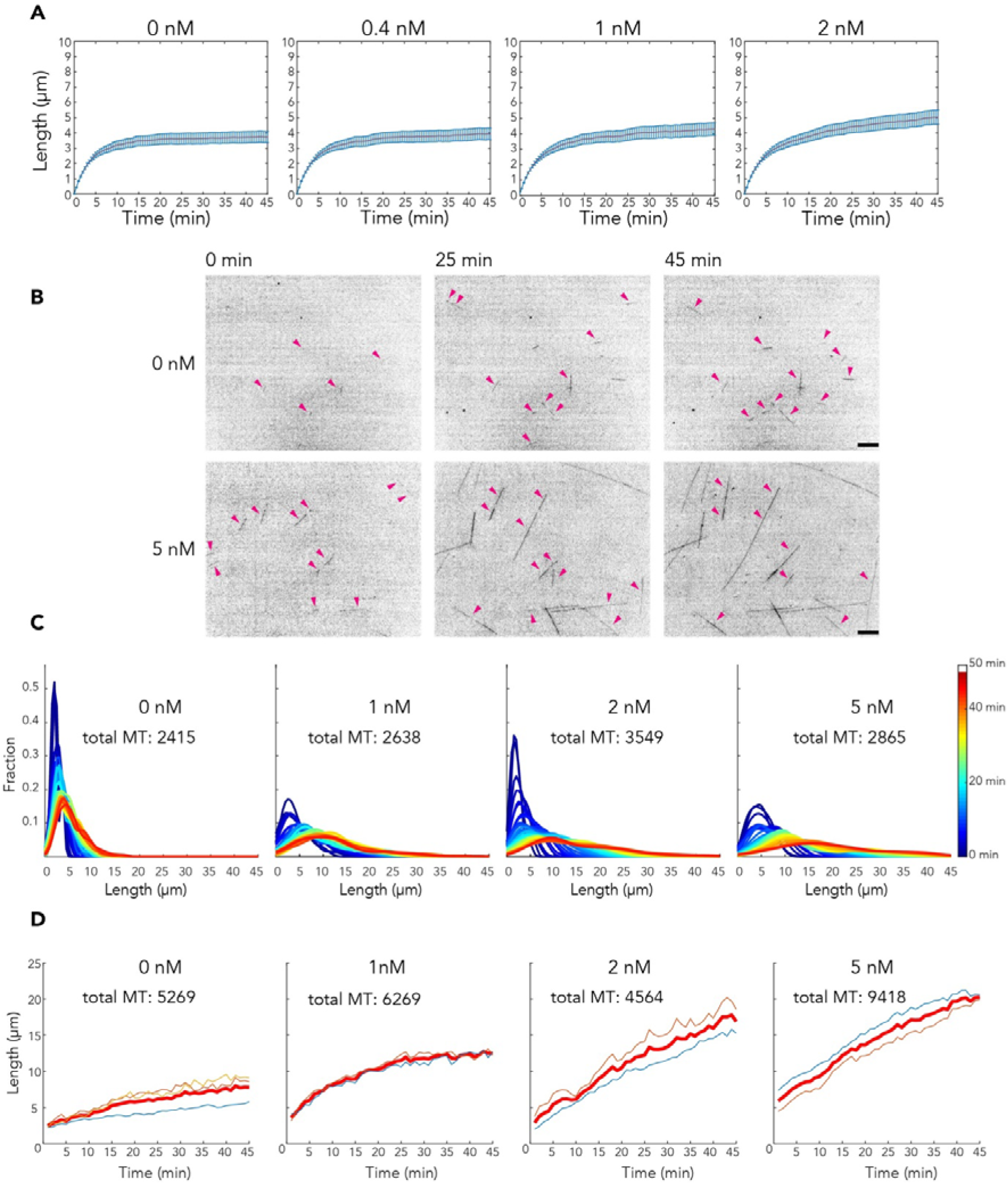
Kinesin-1 controls microtubule length. **(A)** Numerical Simulations: Polymer length versus time for different motor particle concentrations: 0 nM (Control), 0.4nM, 1 nM and 2 nM motor particles. Mean length (red) with SE (blue). Increasing the polymerisation rate of the polymer did not affect the median polymer length (Figure S4D). **(B)** Images from a time lapse showing microtubules (8 µM tubulin) in presence of 0 nM or 5 nM kinesin-1 (imaging rate: 1/min, Video S4 and S5). Addition of kinesin-1 increased the net polymer mass after 25 min in the field of view. The net polymer mass further increased after 45 min. Magenta arrows indicate microtubules which were measured for the statistics in Figures C and D (crossing microtubules and microtubule bundles were excluded from the statistics). Scale bar 10 µm. **(C)** Microtubule length distribution over time (see LUT) at different kinesin-1 concentrations: 0 nM (Control), 1, 2 and 5 nM kinesin-1. Microtubules were imaged every minute for 45 min. To exclude rescue events influenced by bundling or crossing between microtubules, we excluded those events. Total number of microtubules analysed in one experiment for each concentration condition is indicated. **(D)** Mean microtubule length over time of two to three independent experiments at different kinesin-1 concentrations: 0 nM (Control), 1, 2 and 5 nM kinesin-1. Microtubules were imaged every minute for 45 min. Thin coloured curves represent mean length of each of the individual experimental repeats (one experiment set of C). Red curves represent the mean length for the three repeats. Total microtubule numbers analysed are indicated.

In our *in vit*ro experiments, we assessed how the microtubule length depends on kinesin-1 concentration (Figure 4B). The microtubule length distribution was unimodal and the average value as well as the standard deviation increased over time (Figure 4C) (Jeune-Smith & Hess, 2010). The length distribution of microtubules was different with and without kinesin-1 and it depended on motor concentration as observed in the simulations. In the presence of 1nM kinesin-1, the average microtubule length increased and reached a plateau after 30 minutes (Figures 4D and S4E). In these conditions, the average length was larger than without motors (12.6 µm *versus* 5.8 µm after 30 min; Figure 4D). In contrast to the bound growth with 1 nM kinesin-1, for 2 nM kinesin-1, the average microtubule length did not plateau: microtubules continued to grow comparable to the simulations. Together, the *in vitro* and theoretical results show that kinesin-1-induced GTP-tubulin incorporation along the microtubule shaft dramatically increases the microtubule rescue frequency and average length. Above a critical concentration of running kinesin-1 on microtubules, the microtubule length distribution switches from a steady state regime to a behaviour of continuous growth.

### In cells, kinesin-1 activity promotes rescue events and controls microtubule lifetime

The results of our *in vitro* studies and numerical simulations indicated a positive feedback between the frequency at which microtubules are used as tracks by kinesins and the stability of those microtubules. We wondered whether this positive feedback changes the cellular organization of the microtubule network depending on kinesin activity. To address this question, we manipulated the concentration and activity of kinesin-1 in cells, and studied their effects on microtubule network dynamics, density and organization.

To reduce the native kinesin-1 pool by 90 %, we performed an siRNA knockdown of kinesin-1 in CRISPR/Cas9 knock-in GFP-Tubulin HeLa cells (Figures 5A and 5B). This lower concentration of kinesin-1 resulted in a 20% reduction of the rescue frequency compared to control conditions (Figure 5C), in line with a previous observation (Daire et al., 2009). The remaining rescue events could be due to various microtubule associated proteins or mechanical constraints (Aher et al., 2018; Al-Bassam et al., 2010; Arnal et al., 2004; Komarova et al., 2002; Lindeboom et al., 2019). Conversely, overexpression of a constitutive active kinesin-1, K560-mCherry, in Ptk2 cells stably expressing GFP-Tubulin increased the rescue frequency by 50%, depending on the level of K560 expression (Figures 5D, 5E, S5A and S5B). Therefore, kinesin-1 contributed considerably to the generation of rescue sites in the cellular microtubule network.

**Figure 5.**
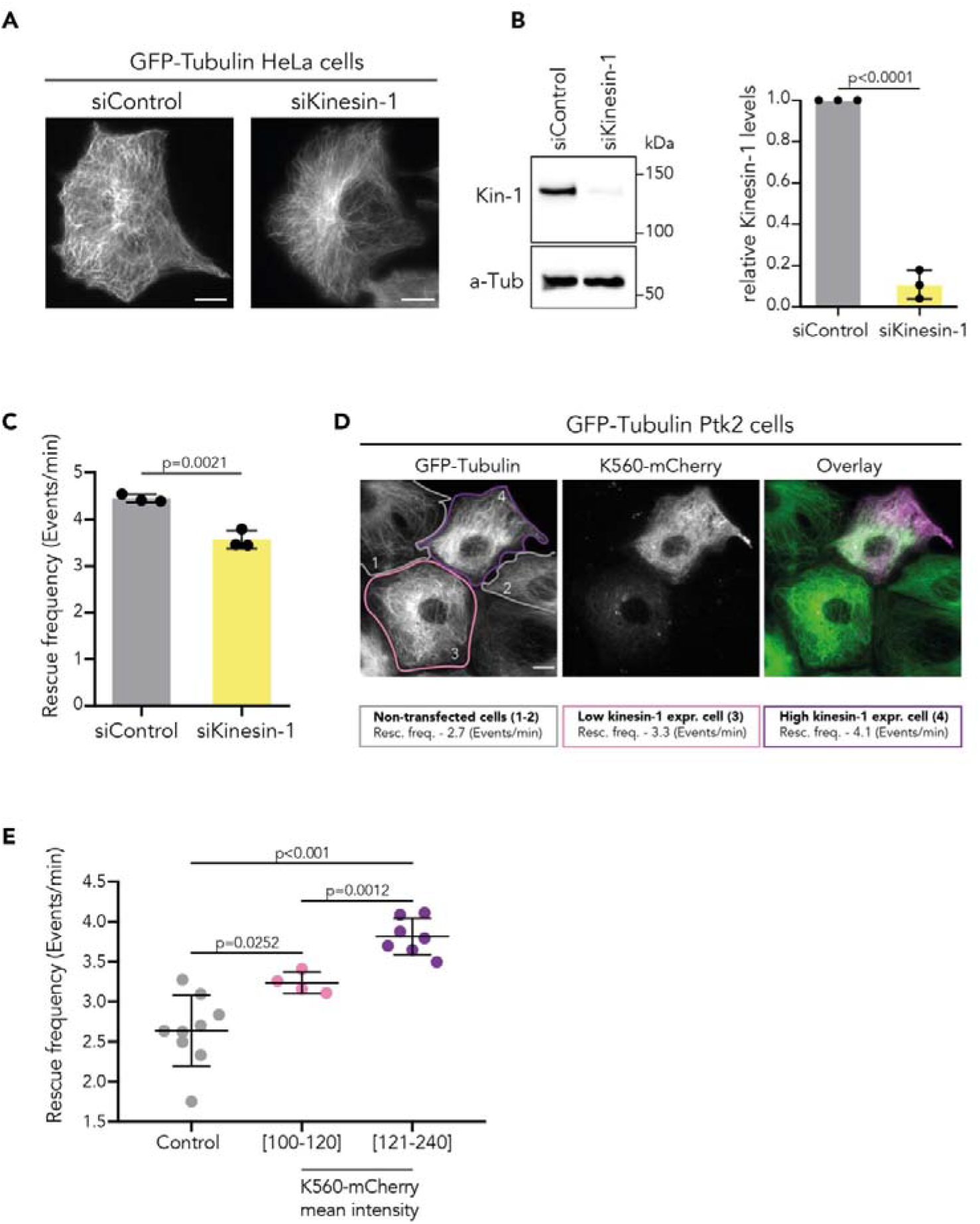
Kinesin-1 activity regulates microtubule rescue frequency in cells. **(A)** Representative images of the microtubule network in siRNA Control (left) and siRNA kinesin-1 (right) GFP-Tubulin HeLa cells. Scale bar: 10 µm. **(B)** Representative Western Blot with quantification of kinesin-1 expression levels in kinesin-1 knockdown GFP-Tubulin HeLa cells. Knockdown of kinesin-1 (siRNA kinesin-1) shows an 80-90% reduction compared to control cells (siRNA Control) from three independent experiments. Statistics: two-tailed T test. Mean with SD. **(C)** Kinesin-1 knockdown in GFP-Tubulin HeLa cells (siRNA kinesin-1) decreases microtubule rescue frequency compared to control cells (siRNA Control). n=200 microtubules from 14 cells (siRNA Control) and n=180 microtubules from 12 cells (siRNA kinesin-1) from three independent experiments. Statistics: two-tailed T test. Mean with SD. **(D)** Representative images of the microtubule network in GFP-Tubulin HeLa cells overexpressing K560-mCherry. Cell 1 and 2 no overexpression, cell 3 low overexpression, cell 4 higher overexpression with the corresponding microtubule rescue frequencies of the cells. Scale bar: 10 µm. **(E)** Rescue frequency measured in GFP-Tubulin HeLa cells overexpressing K560-mCherry at low (n=95 MTs from 4 cells) and higher intensity values (n=150 MTs from 7 cells) compared to non-transfected cells (n=90 MTs from 9 cells) from three independent experiments (see methods). Statistics: unpaired T test. Mean with SD. Representative microtubules analysed are shown in Figure S5.

To study further the effect of kinesin-1 on local microtubule shaft dynamics, we acutely activated native kinesin-1 by using the small molecule Kinesore in our Ptk2 GFP-Tubulin cell line (Figure 6A). The small molecule Kinesore activates the native autoinhibited kinesin-1 pool by interacting with the cargo binding site of the kinesin-1 light chain (Randall et al., 2017). It thereby allows the binding and running of kinesin-1 on microtubules even in the absence of cargo upon addition of the compound. Adding 100 µM Kinesore to Ptk2 cells increased the binding of kinesin-1 to microtubules by a factor of four (Figures 6B, S6A, S6B, S6D and S6E). As a consequence, the microtubule rescue frequency increased by 28% in Ptk2 cells (Figures 6C, 6D, and S6C). This higher rescue rate per microtubule led to a 1.5-fold increase of microtubule lifetime (Figures 4E and 4F). Summing up, increasing the expression level of active kinesin-1 and activating the cellular kinesin-1 pool both resulted in higher microtubule shaft dynamics, higher rescue frequency, and higher microtubule stability, similar to our observation *in vitro* at higher kinesin-1 concentrations (Figure 2D).

**Figure 6.**
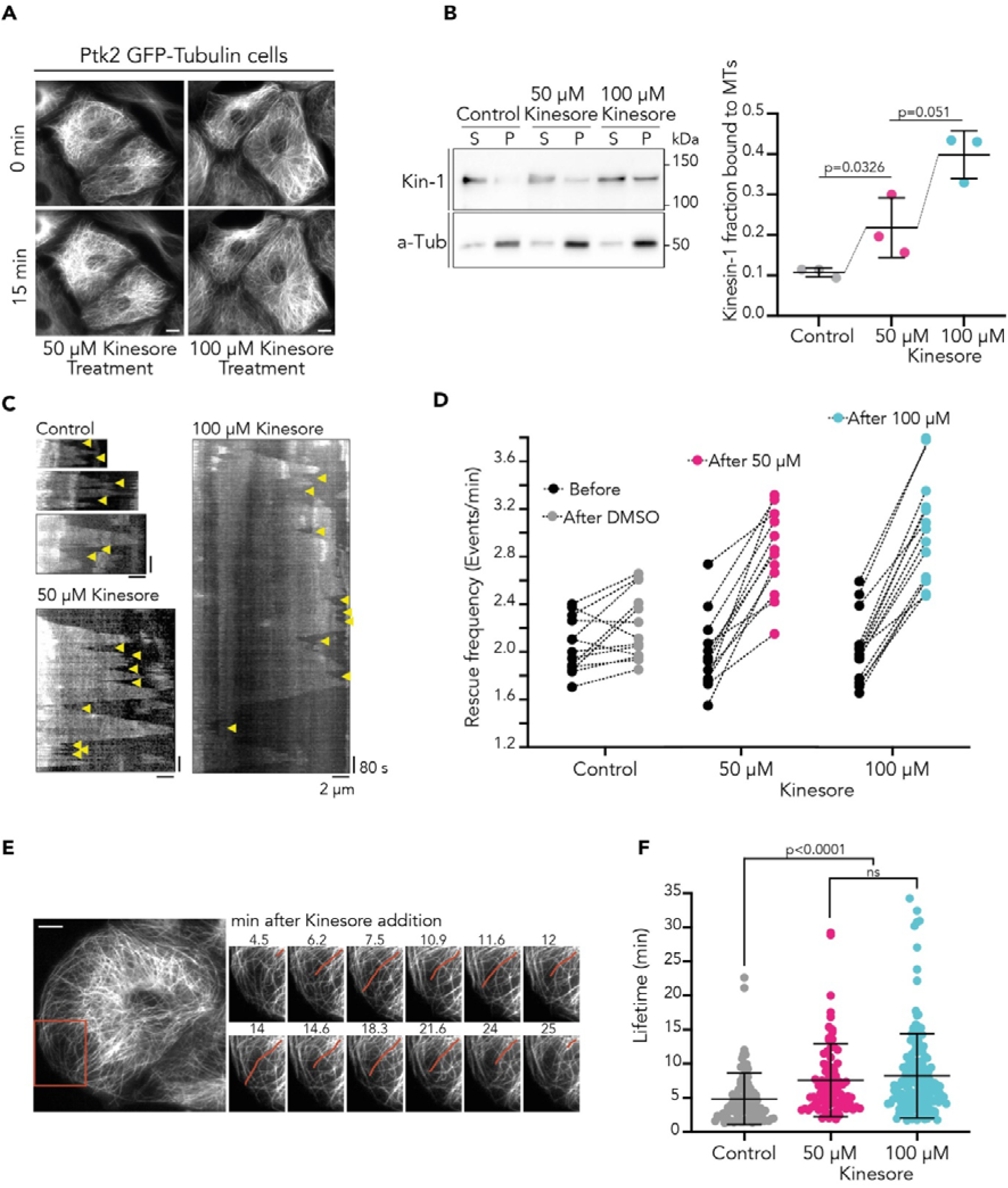
Kinesin-1 activity regulates microtubule dynamics and lifetime in cells. **(A)** Representative images of the microtubule network reorganization in GFP-Tubulin Ptk2 cells before (top) and after 50 µM and 100 µM Kinesore treatment (bottom). **(B)** Kinesore addition for 15 min increases kinesin-1 binding to microtubules in Ptk2 cells (MT, Pellet fraction, P). Representative Western Blot with quantification. Fraction of active kinesin-1 bound to microtubules determined by the Kinesore-dependent microtubule binding assays for kinesin-1 (see methods). Control (0.2 % DMSO) and 100 µM Kinesore treated Ptk2 cells. Mean with SD from three independent experiments. Relative binding values are shown in Figure S6B. Statistics: unpaired T test. **(C)** Representative kymographs of microtubules in GFP-Tubulin Ptk2 cells treated with 0.2 % DMSO, 50 µM Kinesore and 100 µM Kinesore. Yellow arrowheads indicate rescue events. **(D)** Rescue frequency measured in GFP-Tubulin Ptk2 cells before and after treatment with 0.2% DMSO (Control), 50 µM Kinesore and 100 µM Kinesore. Each black circle (to the left) represents the rescue frequency of 10 microtubules in Ringer’s buffer and it is connected to a grey, magenta or cyan circle (to the right) representing the rescue frequency of 10 microtubules after DMSO (n=13 cells), 50 µM (n=12 cells) or 100 µM Kinesore (n=14 cells) is added to the same cells. Fold change in rescue frequency is shown in Figure S6C. **(E)** Representative time-lapse of a microtubule lifetime (highlighted in red) after 100 µM Kinesore treatment in GFP-Tubulin Ptk2 cells. **(F)** Quantification of microtubule lifetime after treatment with 0.2% DMSO (Control; n=90, from 9 cell), 50 µM Kinesore (n=80, from 8 cell) and 100 µM Kinesore (n=156, from 7 cells) in GFP-Tubulin Ptk2 cells from three independent experiments. Statistics: two-tailed T test. Mean with SD. Scale bar: 10 µm.

### Control of rescue by kinesin-1 densifies the microtubule network and rearranges it in space

The kinesin-1 caused increase of microtubule lifetime influences the global microtubule network (Figure 7). We measured the GFP-Tubulin fluorescence intensity distribution in our HeLa cells before and after Kinesore treatment. We segmented the pattern of GFP fluorescence intensity into three categories: low, intermediate and high. We calibrated the intensity in the categories so that they correspond to i) the intensity of cytoplasmic tubulin-dimers (low), ii) the intensity from single microtubules (in locations where individual microtubules can be distinguished; intermediate), and the intensity from higher density networks, where single microtubules cannot be resolved (high; Figure 7B and for details see S7A). Upon Kinesore incubation, the total GFP-Tubulin fluorescence intensity associated to the high-density microtubule network increased at the expense of free tubulin-dimers and isolated single microtubules (Figures 7A, 7B, S7B-E). Note that higher density networks are not due to microtubule bundling, as we verified by expansion microscopy (Figures S8A and S8B). We confirmed the effect of Kinesore treatment on the reduction of the abundance of cytoplasmic tubulin and on the increase of the microtubule network mass by Western blot (Figure S8C). This effect was specific for kinesin-1, since Kinesore treatment did not affect the microtubule network in kinesin-1 siRNA knockdown cells (Figure S7F).

**Figure 7.**
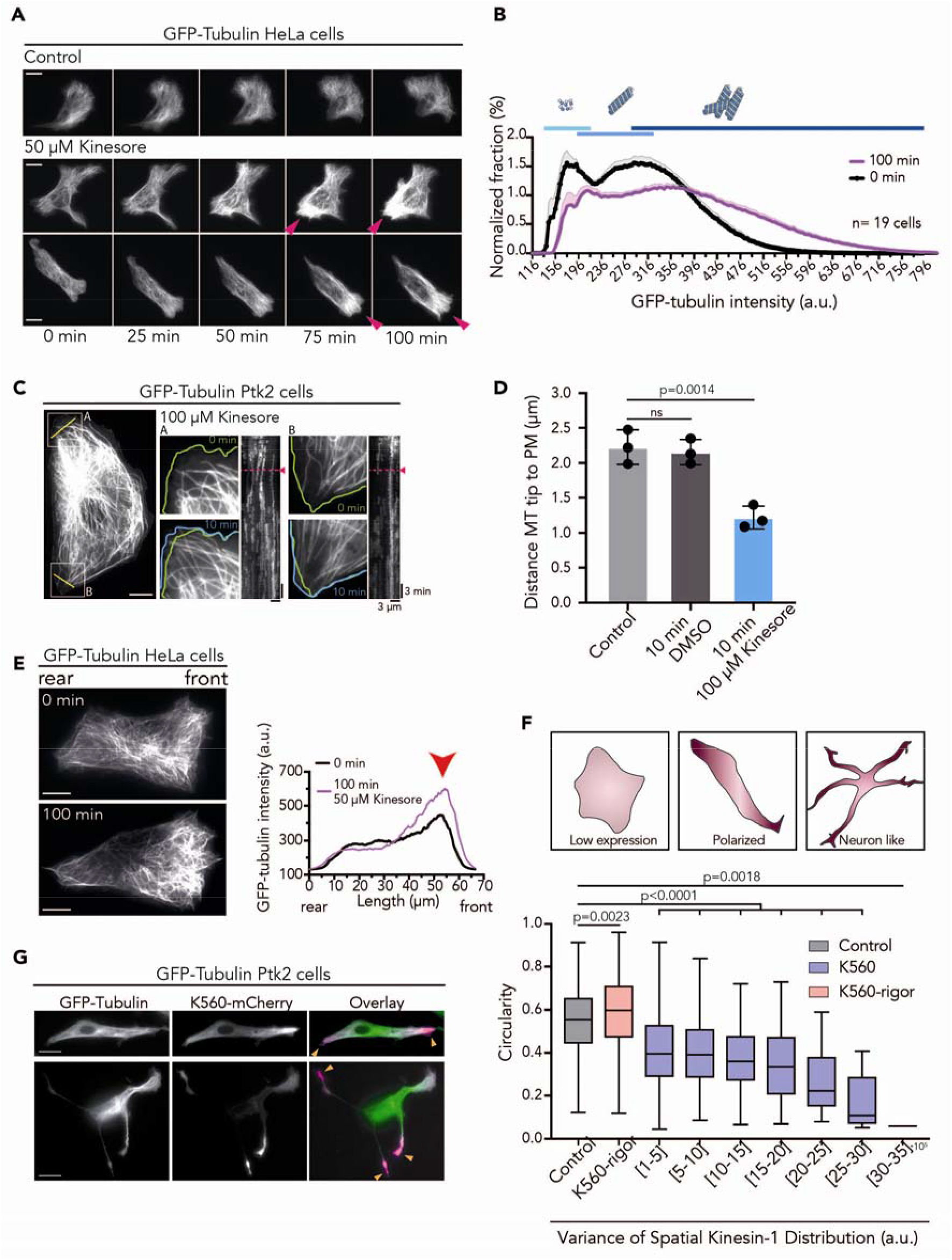
Kinesin-1 regulates microtubule network mass and organization in cells. **(A)** GFP-Tubulin HeLa cells imaged every 5 min for 100 min after 0.1% DMSO (Control) or 50 µM Kinesore addition. The microtubule network organization is not affected by DMSO addition. Addition of Kinesore leads to reorganization of the microtubule network towards the cell rear. The network density increases (magenta arrowheads) where pre-established polarity occurred. **(B)** Distribution of GFP-Tubulin fluorescence intensity before and after 100 min of 50 µM Kinesore treatment, measured in GFP-Tubulin HeLa cells. Mean percentage of total cellular fluorescence GFP-Tubulin intensity (n=19 cells); shaded area indicates SEM. Fluorescence intensity is grouped into three categories, cytosolic dimers (light blue), single microtubules (medium blue), and higher density network (dark blue). Kinesin-1 activation by Kinesore shifts the histogram to higher fluorescence intensities indicating association of tubulin to dense networks of microtubules. For important control conditions see Figure S7. **(C)** Representative GFP-Tubulin Ptk2 cell before and after 10 min of 100 µM Kinesore treatment. The zoom-ins (white squares) show representative microtubule network reorganization upon Kinesore treatment. Kymographs (from yellow lines) show an increase in microtubule number after 4 min of Kinesore addition (magenta arrowhead, dotted line). The effect is enhanced over time (Video S6). Further example shown in Figure S8D. **(D)** Distance from the microtubule (MT) tips to the plasma membrane (PM) before treatment (Control), and after 10 min treatment with 0.2% DMSO, or 100 µM Kinesore. For each condition three independent experiments were performed: Control; n=90 MT from 12 cells, 0.2% DMSO; n=80MT from 12 cells, 100 µM Kinesore; n=156 MT from 12 cells). Statistics: two-tailed T test. Mean with SD. **(E)** GFP-Tubulin HeLa cell before and after 100 min incubation with 50 µM Kinesore, showing the amplification of a pre-existing heterogeneity in density of the microtubule network following Kinesore treatment (red arrow). **(F)** Cell circularity of non-transfected stable expressing GFP-Tubulin Ptk2 cells compared to GFP-Tubulin Ptk2 cells overexpressing K560-rigor and K560. Cell circularity is plotted over binned variance of spatial K560 distribution. Red colours in the scheme indicate kinesin-1 concentration. Statistics: one-way ANOVA. Boxplots show median with the 25th and 75th percentile as box edges. Control (non-transfected cells, n=26 cells), K560-rigor (n=58 cells) and K560 (n=67 cells) from three independent experiments. Changes in circularity over increasing mean fluorescence intensity values of K560 and K560-rigor are shown in Figure S9C. **(G)** Asymmetric K560-mCherry distribution impacts on the morphology of GFP-Tubulin Ptk2 cells. Orange arrowheads highlight regions of higher kinesin-1 intensity. Scale bars: 10 µm.

The denser microtubule network after kinesin-1 activation was particularly apparent at the cell periphery (Figures 7A, 7C and S8D), where microtubules were sparse in Ptk2 control cells (Provance et al., 1993). Indeed, the average distance between the plasma membrane and the microtubule tip was reduced by half after 10 min of Kinesore treatment (Figure 7D, Video S6 and S7). In particular pre-existing heterogeneities within the microtubule network get amplified upon Kinesore activation of kinesin-1 (Figures 7E). To increase kinesin-1 cargo-dependent transport we overexpressed SKIP, a lysosomal adaptor protein, in our Ptk2 GFP-Tubulin cells. Increased lysosomal transport densified the peripheral microtubule network, in-line with our observation for running kinesin-1 without load (Figures S9A and S9D). Therefore, both running kinesin-1 without and with cargo densify the peripheral microtubule network.

The microtubule network determines cell polarity. By modulating shaft dynamics, kinesin-1 could thus affect cell morphology. Indeed, cell circularity decreased monotonously up to four-fold with increasing expression levels of K560-mCherry (Figures S9B and S9C). Cell circularity decreased even stronger – up to nine-fold – with the variance of kinesin-1 spatial distribution (Figures 7F and 7G, Video S8). At high variance of K560-mCherry spatial distribution we observed cells with neuron-like morphologies (Figure 7G). This anisotropic cell morphology was also caused by increased kinesin-1 mediated lysosomal transport due to SKIP overexpression (Figure S9D). Conversely, overexpression of a rigor K560-mCherry mutant did not reduce, but actually increased the circularity of our Ptk2 cells (Figures 7F, S9B and S9C). Concluding, only active kinesin-1 can break cell symmetry by controlling microtubule dynamics at the shaft.

## DISCUSSION

Taken together, our data show that kinesin-1 is not only using the microtubules as tracks for cargo delivery in cells, but, at the same time, it controls the length and lifetime of their preferred tracks. This is based on the following observations: i) running kinesin-1 increases microtubule rescue events (Figure 2D), ii) average length of microtubules increases depending on how frequently they are used as motor tracks (Figure 4), iii) active kinesin-1 densifies the cellular microtubule network and this enhances cell polarity (Figures 6 and 7).

The underlying principle of running kinesin-1 modulating microtubule dynamics, is to causes tubulin dissociation from the shaft. One might assume that kinesine-1 is pulling out a dimers, but this is not the case as our experimental data showed no tight coupling between the step of a kinesin-1 molecule and the removal of a tubulin dimer from the shaft. Indeed, only a small fraction of kinesin-1 steps led to tubulin removal. Also binding of kinesin-1 does not increase tubulin dissociation, as our data for immotile kinesin-1 show (Figure S3D). Instead, experiments with no kinesin-1 show that spontaneous tubulin dissociation from the shaft is a stochastic process driven by thermal energy (Schaedel et al., 2019). We conclude that running kinesin-1 increases the probability of thermal tubulin dissociation (Figure 1).

Kramer’s rate theory shows that the thermal tubulin dissociation rate scales like exp(-ΔE/kT), where ΔE is the height of the energy barrier that has to be overcome for dissociation and kT is the thermal energy (k: Boltzmann constant, T: temperature) (Risken, 1984). Reducing the energy barrier for a tubulin dimer to leave the shaft by less than 1 kT already increase the dissociation rate by a factor of 2. The energy derived from ATP hydrolysis under cellular conditions is ≈ 25 kT (Jonathon Howard, 2001). The maximal work of kinesin-1 under load is 48 pN*nm, requiring ≈ 10 kT per step (Schnitzer & Block, 1997). Therefore energy from ATP hydrolysis is sufficient to increase the probability of spontaneous tubulin dissociation from the shaft, even when the motor is under load.

In the scenario where kinesin-1 controls microtubule stability by rescue, a positive feedback loop ensures that frequently used tracks become more stable. It is well established that kinesin-1 shows preferential binding to a subset of long-lived microtubules which are also acetylated and/or detyrosinated (Cai et al., 2009; Hammond et al., 2010). However *in vitro* experiments showed only slight effects on the affinity of kinesin-1 for tubulin with posttranslational modifications (Kaul et al., 2014; Soppina et al., 2012). It is therefore unclear whether preferential binding to microtubules with these post-translational modifications is a cause or a consequence of these modification. Our observations are indicating that kinesin-1 controls the lifetime of a subset of microtubules which thereafter get post-translationally modified due to their increased lifetime.

Beyond the microtubule tips, including the microtubule shaft as a regulatory structure changes our understanding of the microtubule network from i) a spotted system which is only controlled in discrete points at the end of filaments to ii) a continuous system controlled and modified everywhere in the network. To support cell polarity, the microtubule network needs to be functionally and structurally asymmetric. The symmetry of a network array can be broken via four general mechanisms: relocation of existing microtubules within a cell; addition of an asymmetric nucleation site; repositioning of nucleation sites homogeneously distributed in the network to one side of a system; differential modulation of microtubule dynamics.

Kinesin-1 can break the network symmetry by differential modulation of microtubule dynamics. The preferential binding of kinesin-1 to a subset of microtubules within the network causes local stabilization and local increase in density of microtubules, which in turn increases kinesin-1 binding in that region. Indeed, this kinesin-1/microtubule positive feedback system can generate from minor heterogeneities in space, as an emerging property, a heterogeneous landscape of microtubules underlying cell polarity. Any slight spatial heterogeneity in the concentration of running kinesins will be then amplified. We did observe such amplifying heterogeneities upon cargo-dependent and independent kinesin-1 activation as well as a consequence of overexpression of kinesin-1 (Figures 7A, 7E-G and S9B, S9D). Therefore, the activity of the running motors can generate polarity inside the cell or amplify network heterogeneities encoded in other polarity cues (Par complex, planar cell polarity, directed migration, etc.).

## Acknowledgments

We thank O. Schaad for support with the statistical analysis, and M. Gonzalez-Gaitan for scientific discussions. J. Schaer for discussions of the numerical simulations. S. Diez for sharing the Kinesin-1 plasmid and Kinesin purification experiences. M. Gonzalez-Gaitan, P. Guichard, V. Hamel, S. Hoogendoorn, and A. Roux for carefully reading of our manuscript. MAC and SF have been supported by the SNSF, 31003A_182473; KK has been supported by the NCCR Chemical Biology program; CA has been supported by the DIP of the Canton of Geneva, SNSF (31003A_182473), and the NCCR Chemical Biology program.

## Author Contributions

MAC performed the experiments with the help of SF. MAC and CA designed the experiments. MCV purified the proteins and generated the CRISPR cell line. MAC, SF, MCV and CA analysed data. KK designed and performed numerical simulations. CA wrote the manuscript.

## METHODS

### Cell culture, transfection, CRISPR knock-in, and siRNA knockdown

HeLa (ATTC, CCL-2™, a kind gift of Jean Gruenberg) cells were cultured in high glucose Dulbecco’s Modified Eagle’s Medium (DMEM, ThermoFisher, 61965026) supplemented with 10 % Fetal Bovine Serum (FBS, ThermoFisher, 10270106) and 1 % penicillin-streptomycin (Gibco, 15140122) at 37°C with 5 % CO_2_. Ptk2 cell line stably expressing GFP-Tubulin (a kind gift of Franck Perez) was grown in Minimum Essential Medium Eagle (MEM, Sigma, M0643) supplemented with 10 % Fetal Bovine Serum, 1 % penicillin-streptomycin and 1 % L-Glutamine (Gibco, 25030024) at 37°C with 5 % CO_2_. The cell lines were monthly checked for mycoplasma contamination. In this study two cell lines were chosen depending on experimental setups. Ptk2 cells are flat with a sparse microtubule network making visualization and quantification of microtubule dynamics easier. Whereas the human HeLa cell line was used to perform siRNA knockdown assays.

GFP-Tubulin Ptk2 cells were transfected with jetOPTIMUS transfection reagent (Polyplus) using 1 µg of plasmid in 24-well plates according to the manufacturer’s instructions. Cells were analysed 8-24 hours after transfection. The constitutively active form of kinesin-1 (human KIF5B, K560) was cloned by PCR from pet17-K560-GFP_His (Addgene, 15219) into pEGFP-N1 vector using EcoRI primer (104-5p EcoRI-K560: 5’-CTTCGAATTCATATGGCGGACCTGGCCGAGTG CAAC-3’) and AgeI primer (105-3p AgeI-K560 : 5’-GGGACCGGTGGACCTTTTAGTAAAGATGCCATCATCTCA-3’). The GFP sequence was removed from K560-GFP and mCherry AgeI/NotI was inserted instead (K560-mCherry). The ATPase rigor mutant of K560 (K560-rigor-mCherry) was generated by inserting a point mutation (G234A) using 5’-CTaGTTGATTTAGCTGcTAGTGAAAAGGTTAGT and 3’-ACTAACCTTTTCACTAgCAGCTAAATCAACtAG primers. The plasmid encoding full length SKIP (myc-SKIP) was a kindly gift from Dr. Juan Bonifacino.

HeLa CRISPR/Cas9 Tubulin-GFP knock-in cells were generated based on a previously described protocol (Koch et al., 2018). A gRNATubulin sequence, GATGCACTCACGCTGCGGGA, was cloned in pX330 and a plasmid donor, TubA1B-mEGFP (Addgene, 87421), was used for cell transfection. HeLa cells at 80 % confluency were transfected with a solution of 1 µL lipofectamine, 1.5 µg pX330gRNATubulin and 2 µg plasmid donor vector in a 12 wells-plate. After 7-10 days, a Fluorescence-activated cell sorting (Biorad S3) was performed with 7×10^6^ transfected cells (6.000 cells sorted expressing GFP). Cells were grown at 37°C with 5 % CO_2_. After 7-10 days, a second FACS (Sony SH800 Cell sorter) was performed with 80 % confluent pre-selected cells (sorting of single cell in a 96 wells-plate). Cell clones were grown at 37°C with 5 % CO_2_. After 13 days of incubation, the selected clone was picked for correct gene insertion (control PCR) in function of its dividing rate (division rate was reduced by 74 % compared to WT) and fluorescent appearance (visualized at the TIRF microscope).

For generating kinesin-1 knockdown cells, a combination of four siRNA duplexes against Kif5B subunit of kinesin-1 (GeneSolution siRNA, Quiagen) at a final concentration of 10 nM were transfected in HeLa GFP-Tubulin CRISPR knock-in cells using Lipofectamine™ RNAiMAX (ThermoFisher) following the manufacturer’s instructions for 72h. AllStars Negative Control siRNA (10 nM) was used as a negative control in all knockdown experiments. To measure the suppression of kinesin-1 expression by siRNA, the siRNA-transfected cells were lysed [50 mM Tris-HCl (pH 7.5), 150 mM NaCl, 1 % Triton X-100, 0.5 % SDS, and a protease inhibitor tablet (Roche)] for immunoblotting analysis (see SDS-PAGE and Western blot section and Figure 4A).

### Tubulin purification from bovine brain and tubulin labelling

Tubulin was purified from fresh bovine brain by two cycle of polymerisation and depolymerisation as previously described (Castoldi & Popov, 2003). A first polymerisation-depolymerisation cycle was performed in High-Molarity PIPES buffer [1 M PIPES-KOH at pH 6.9, 10 mM MgCl_2_, 20 mM EGTA, 1.5 mM ATP and 0.5 mM GTP] supplemented with 1:1 glycerol and Depolymerisation buffer [50 mM MES-HCl at pH 6.6 and 1 mM CaCl_2_] respectively. A second polymerisation-depolymerisation cycle was then performed: polymerisation in High-Molarity PIPES buffer and depolymerisation in 0.25XBRB80 complete after 15 min with 5XBRB80 to reach 1XBRB80 [80 mM PIPES at pH 6.8, 1 mM MgCl2 and 1 mM EGTA] respectively.

Labelled tubulin with ATTO-488, ATTO-565, or ATTO-647 (ATTO-TEC GmbH) and biotinylated tubulin were prepared as previously described (Hyman et al., 1991) with slight modification. Tubulin was polymerised in Glycerol PB solution [80 mM PIPES-KOH at pH 6.8, 5 mM MgCl_2_, 1 mM EGTA, 1 mM GTP and 33 % (v/v) glycerol] for 30 min at 37°C and layered onto cushions of 0.1 M NaHEPES at pH 8.6, 1 mM MgCl_2_, 1 mM EGTA and 60 % (v/v) glycerol followed by centrifugation. Pellet was resuspended in Resuspension buffer [0.1 M NaHEPES at pH 8.6, 1 mM MgCl_2_, 1 mM EGTA, 40% (v/v) glycerol] and incubated 10 min at 37°C with 1/10 volume of 100 mM ATTO-488, -565, or -647 NHS-fluorochrome or incubated 20 min at 37°C with 2 mM Biotin reagent. Labelled tubulin was sedimented onto cushions of BRB80 supplemented with 60 % glycerol, resuspended in BRB80, and a second polymerisation-depolymerisation cycle was performed before use. Labelling ratio was 11 % for ATTO-488 and 13 % for ATTO-565.

### K430-GFP expression and purification

All *in vitro* experiments were performed with truncated, EGFP-labelled kinesin-1 construct, K430-EGFP (rat KIF5C, referred as kinesin-1 or K430 in the text). The K430-GFP plasmid (a kind gift of Stefan Diez’s laboratory) was expressed and purified as previously described (Rogers et al., 2001). *E*.*coli* BL21(DE3)[pLysS] expressing K430-GFP were lysed in a Lysis buffer [50 mM Na-Phosphate buffer at pH 7.5, 300 mM KCl, 10 % Glycerol, 1 mM MgCl_2_, 20 mM ß-Mercaptoethanol, 0.2 mM ATP, 30 mM imidazole and protease inhibitors cocktail tablets (Roche)]. The cleared lysate was loaded into a pre-equilibrated HisTrap column (GE Healthcare 1mL HisTrap column) using an ÄKTA Pure Protein Purification System (GE Healthcare). After washes, protein was eluted with Elution buffer [50 mM Na-Phosphate buffer at pH 7.5, 300 mM KCl, 10 % Glycerol, 1 mM MgCl_2_, 0.2 mM ATP, 300 mM imidazole and 10 % (w/v) sucrose]. A dialysis was performed overnight to exchange the Elution buffer 20 mM NaHEPES at pH 7.7, 150 mM KCl, 1 mM MgCl_2_, 0.05 mM ATP, 1 mM DTT and 20 % (w/v) Sucrose. Protein concentration was measured by Bradford method and the concentration was adjusted to 1.2 µg/µL using Centrifugal filter Amicon 30K (Millipore). Protein was aliquoted and stored in liquid nitrogen.

### Imaging

Microscopy imaging was realized with an Axio Observer Inverted TIRF microscope (Zeiss, 3i) and a Prime BSI (Photometrics). A 63X objective (Plan-Apochromat 63X/1.46 Oil Korr TIRF, Zeiss) and a 100X objective (Zeiss, Plan-Apochromat 100X/1.46 oil DIC (UV) VIS-IR) were used. SlideBook 6 X64 software (version 6.0.17) was used to record time-lapse imaging. For *in vitro* and cell imaging, microscope stage conditions were controlled with the Chamlide Live Cell Instrument incubator (37°C and 5 % CO_2_).

### Flow chamber

Slides and coverslips were cleaned by two successive incubations and sonication: sonicated for 40 min in 1 M NaOH, rinsed in bidistilled water, sonicated in ethanol (96 %) for 30 min and rinsed in bidistilled water. Slides and coverslips were dried with an air gun, placed into a Plasma cleaner (Electronic Diener, Plasma surface technology) for plasma treatment, followed by 2 days incubation with tri-ethoxy-silane-PEG (Creative PEGWorks) or a 1:5 mix of tri-ethoxy-silane-PEG-biotin and tri-ethoxy-silane-PEG at 1 mg/ml in 96 % ethanol and 0.02 % HCl, with gentle agitation at room temperature. Slides and coverslips were then washed in ethanol (96 %), and bidistilled water, dried with air gun and stored at 4°C. Flow chamber was assembled by fixing with double tap a tri-ethoxy-silane-PEG treated slide with a 1:5 mix of tri-ethoxy-silane-PEG-biotin and tri-ethoxy-silane-PEG treated coverslip.

### Incorporation experiment in presence of ATP and kinesin-1

Microtubule seeds were prepared at 10 µM tubulin concentration (20 % ATTO-647-labelled tubulin and 80 % biotinylated tubulin) in BRB80 supplemented with 0.5 mM GMPCPP (Jena Bioscience) for 1 hour at 37°C. Seeds were incubated with 1 µM Paclitaxel (Sigma) for 45 min at 37°C, centrifuged (50.000 rpm at 25°C for 15 min), resuspended in BRB80 supplemented with 1 µM Paclitaxel and 0.5 mM GMPCPP and stored in liquid nitrogen.

Flow chamber was prepared by injecting successively 50 µg/mL neutravidin (ThermoFisher), BRB80, GMPCPP microtubule seeds, and washed with BRB80 to remove unattached seeds. According to Figure S1A, microtubule polymerised from the seeds with 8 µM tubulin (20 % Atto-488 labelled) in BRB80 supplemented with an anti-bleaching buffer [10 mM DTT, 0.3 mg/mL glucose, 0.1 mg/mL glucose oxidase, 0.02 mg/mL catalase, 0.125 % methyl cellulose (1500 cP, Sigma), 1 mM GTP and 2.7 mM ATP] and 0.2% BSA. The chamber was incubated for 15 min at 37°C for polymerisation. To cap microtubules, the polymerisation buffer was exchanged with 5 µM tubulin (70 % Atto-488 labelled) in BRB80 supplemented with 0.5 mM GMPCPP and incubated for 12 min at 37°C. An incorporation buffer containing 9 µM tubulin (100 % Atto-565 labelled) and 0, 1, 2 or 5nM K430-GFP in BRB80 supplemented with anti-bleaching buffer, 0.2% BSA and 41 mM KCl was flushed-in the chamber and incubated for 15 min at 37°C. A tubulin-free BRB80 containing 0.2% BSA wash was performed and incubated for 1 min at 37°C to depolymerise all dynamic tubulin structures grown on top of existing microtubules or beyond the stabilizing GMPCPP cap. 6 µM tubulin (100 % black) in BRB80 with anti-bleaching buffer and 0.2% BSA was injected. The chamber was sealed for microscopy. Note that the concentration of free tubulin did essentially not change in the course of an experiment. We recorded each position over 10 frames and only analysed microtubules not sticking to the cover glass, to ensure that measured signal was an incorporation site and not an aggregate on the surface, (see Figure 1A). To measure incorporation sites, we first calculated the fluorescence intensity of one protofilament. We therefore performed a line-scan (3-pixel wide line with ImageJ) along microtubule fractions growing beyond the GMPCPP-cap, these microtubules are polymerized from 100% Atto-565 labeled tubulin. We subtracted the background from the signal and divided it by 14, the majority of microtubules grown from GMPCPP have 14 protofilaments (Vemu et al., 2017). We defined an incorporation site as a fluorescence signal equal to one or more protofilaments. We counted incorporations separately when the signal dropped below one protofilament in-between two incorporation sites.

To measure the integrated fluorescence intensity of incorporation sites we performed a line-scan (4-pixel wide line with ImageJ) along incorporation sites. We measured the background fluorescence intensity exactly the same position with a second line-scan, therefore we choose a time frame where the microtubule with the incorporation site fluctuated out of field of measurement. To retrieve the integrated fluorescence intensity of exchanged dimers within the incorporation, we subtracted the background raw intensity from the incorporation raw intensity.

For kinesin-1 velocity measurement, the GDP-microtubule were polymerised and capped as described above, followed by injecting 0.4 nM K430-GFP, 8 µM tubulin with anti-bleaching buffer, 0.2% BSA and 41 mM KCl. Images were recorded every second with a laser power of 2.5 mW for 488 nm and 1.3 µW for 561 nm to minimize photobleaching. Kinesin-1 velocity was measured from kymographs build with MultipleKymograph in ImageJ.

### Microtubule dynamics and length in presence of ATP and kinesin-1

Microtubules were polymerised from microtubule seeds (see Incorporation experiment above) with 8 µM tubulin (20 % Atto-565 labelled) in BRB80 supplemented with anti-bleaching buffer, 0.2% BSA and 41.6 mM KCl and incubated for 10 min at 37°C. Then, BRB80 supplemented with anti-bleaching buffer, 0.2% BSA and 41.6 mM KCl containing 0, 0.4, 1, 2, or 5 nM K430-GFP was injected and the chamber sealed for imaging of microtubule dynamics. Images were recorded every 3-4 sec with the lowest laser power possible for our system 1.3 µW. To further reduce phototoxicity we imaged K430-GFP (2.5 mW for 488 nm) only every 30 frames. Polymerisation and depolymerisation speed, catastrophe and rescue frequency were measured from kymographs build with MultipleKymograph in ImageJ. For control conditions to assure kinesin-1 motility we imaged kinesin-1 together with microtubules every 1 or 3 sec, to minimise the phototoxicity in control conditions we imaged just microtubules.

For microtubule length measurement, a solution of 8 µM tubulin (20 % labelled) in BRB80 supplemented with an anti-bleaching buffer, 0.2 % BSA, 41.6 mM KCl and 0, 0.4, 1, 2, or 5 nM was perfused inside the chamber, the chamber sealed and directly imaged at the microscope. To reduce as much as possible any phototoxicity effect we imaged only the microtubule channel, and images were recorded every min for 45 min with the same laser power as before. Microtubule length was measured manually using ImageJ. We did not differentiate between minus and plus end growth. The length distribution and the length over time were analysed with MATLAB.

### Microtubule dynamics in presence of AMP-PNP and kinesin-1

For AMP-PNP experiment, 6.5 µM tubulin was used (instead of 8 µM) to have dynamic microtubules in the presence of higher MgCl_2_ concentrations (100 mM AMP-PNP is dissolved in 100 mM MgCl_2_ solution and not in H_2_O like ATP). An elongation solution of 6.5 µM tubulin (20 % labelled) in BRB80 supplemented with 0.2 % BSA and anti-bleaching buffer (10 mM DTT, 0.3 mg/mL glucose, 0.1 mg/mL glucose oxidase, 0.02 mg/mL catalase, 0.125 % methyl cellulose (1500 cP, Sigma), 1 mM GTP, 2.7 mM MgCl_2_ and 2.7 mM AMP-PMP), 80 mM KCl and 2 nM K430-GFP (or 0 nM for Control) was injected inside the flow chamber. In the flow chamber the microtubules polymerised from GMPCPP microtubule seeds. Images were recorded every 3-4 sec with a laser power of 2.5 mW for 488 nm and 1.3 µW for 561 nm. To reduce phototoxicity on microtubule we imaged K430-GFP (2.5 mW for 488 nm) only every 30 frames. Depolymerisation speed, catastrophe and rescue frequency were measured from kymographs build with MultipleKymograph in ImageJ. As a control, kinesin-1 binding to microtubules was imaged together with microtubules every 3 sec.

### Kinesore treatment

Kinesore treatment was adapted from a previously published method (Randall et al., 2017). The small molecule Kinesore was solubilized in DMSO at a concentration of 50 mM, aliquoted and stored at -20°C. All Kinesore treatment experiments were performed in Ringer’s buffer: 150 mM NaCl, 5 mM KCl, 2 mM CaCl_2_, 2 mM MgCl_2_, 10 mM HEPES, 11 mM glucose pH 7.4 without CO_2_, at 37°C. Our Ringer’s buffer is slightly modified from the buffer used by Randall et al. which increase the viability of our cells from under 2 hrs to over 6 hrs, and abolished the described reorganisation of the microtubule network into loops and thick bundles in our hands (Randall et al., 2017). In control experiments DMSO was added at 0.1 % (control for 50 µM Kinesore) or 0.2 % (control for 100 µM Kinesore).

### Live-cell Imaging

GFP-Tubulin CRISPR knock-in HeLa cells and stable expressing GFP-Tubulin Ptk2 cells (kind gift from Franck Perez) were grown on glass coverslips for microscopy live cell imaging. To reduce photobleaching the laser power was controlled to 0.86 mW, for 488 nm. Videos were processed using ImageJ and linear adjustments (brightness and contrast) were performed.

For <u>microtubule dynamic measurements</u> images were taken every 5 sec. Individual microtubules were manually tracked over time with the Freehand lines tool (see Figure S5B) and the corresponding kymographs were drawn using KymographBuilder in ImageJ. Rescue frequency was measured and calculated from kymographs by dividing the total rescue events by the total depolymerisation time. Complete depolymerisation (end of microtubule life) was considered when the microtubule could not be followed anymore in the field of view.

The effect of reduced kinesin-1 expression on microtubule dynamic instability was analysed in kinesin-1 knockdown GFP-Tubulin HeLa cells (siKinesin-1, Quiagen) as well in control cells (siControl, Quiagen). For details see Cell culture section above. A minimum of 10 microtubules were analysed per cell and 3-4 cells per experiment.

The effect of constitutive active kinesin-1 (K560-mCherry) on microtubule rescue frequency was measured in GFP-Tubulin Ptk2 cells after 16 h of overexpression. We assigned the K560-mCherry overexpressing cells into two categories dependent on the mean cellular grey value of the mCherry channel: low K560-mCherry expression levels (cells with a mean grey value of 100 to 120) and higher expression levels (cells with a mean grey value of 120 to 240). In order to calibrate and compare mean intensities values from independent experiments Fluorescent beads (TetraSpeck™ Microspheres, 1.0 µm; T7282, ThermoFisher) were used. Examples of microtubules selected for analysis are shown in Figure S5B.

The effect of Kinesore induced kinesin-1 activation on microtubule rescue frequency was observed in GFP-Tubulin Ptk2 cells. After imaging cells every 5 sec for 10 min in Ringer’s buffer, 0.2 % DMSO, 50 µM Kinesore, or 100 µM Kinesore were added. We continued imaging the cells every 5 sec for further 40 min at 37°C. A minimum of 10 microtubules were analysed per cell before and after treatment and 3-4 cells per experiment. To determine the lifetime (min) of dynamic microtubules the same kymographs were used.

To <u>measure changes in the distance from the microtubule tip to the plasma membrane</u> after Kinesore treatment, images of GFP-Tubulin Ptk2 cells were analysed at timepoint 0 min and 10 min after treatment with 0.2 % DMSO (Control) and 100 µM Kinesore. For analysis two regions per cell were chosen and all microtubule perpendicular to the plasma membrane were analysed within the region. The minimal distance between single microtubules and the plasma membrane was measured using the line tool from ImageJ.

<u>Changes in microtubule network density</u> upon Kinesore treatment was studied in GFP-Tubulin HeLa cells, and kinesin-1 knockdown GFP-Tubulin HeLa cells (siKinesin-1). We measured the GFP-Tubulin fluorescence intensity distribution, for detail about calibration of the intensities see Figure S7A. Images were taken every 5 min for 100 min, one before and 19 images after addition of 0.1 % DMSO (Control), or 50 µM Kinesore. The cellular GFP-Tubulin fluorescence intensity distribution was measured in ImageJ using Histogram with a bin of 4. The fraction of fluorescence intensity was normalized to each timeframe. In the histogram of fluorescence intensity, the mean normalized fraction of all cells measured per experiment was plotted (n=16 to 24 cells) (Figures 7B and SB-F).

<u>Kinesin-1 impacting cell morphology</u> was studied by measuring cell circularity of non-transfected Ptk2 cells stable expressing GFP-Tubulin and cells overexpressing K560-mCherry and K560-rigor-mCherry. Starting 16 h after transfection we imaged every hour for 48 h. Circularity (4π*[area/perimeter^2^]) was calculated using ImageJ, using the cell contour of single cells every timeframe. Both mean fluorescence intensity and the variance of kinesin-1 spatial distribution changes over time were measured and binned in 8 and 7 groups (Figures S9C and 7F, respectively) and plotted over cell circularity.

To analyse whether <u>cargo-dependent transport</u> impacts on microtubule network reorganization and cell shape, Ptk2 GFP-Tubulin cells were transfected with myc-SKIP (lysosomal adaptor protein) to increase kinesin-1 mediated lysosome transport in the cells (Guardia et al., 2016). To follow lysosome movement in live cells, Ptk2 cells were incubated with Lysotracker (1:15000 dilution; 67 nM) for 10 min at 37°C before imaging.

### Tubulin extraction assay

The tubulin extraction assay was modified from a previously described method (Gasic et al., 2019).

HeLa or Ptk2 cells were grown until 80-90 % confluency in 12-wells plates and treated with 0.1 % DMSO (Control HeLa), 0.2 % DMSO (Control Ptk2), 50 µM, or 100 µM Kinesore in Ringer’s buffer for 40 min at 37°C. Cells were rinsed once with warm PBS before lysing them with 300 µl warm tubulin extraction buffer [60 mM PIPES at pH 6.8, 25 mM HEPES (pH 7.2), 10 mM EGTA at pH 7-8, 2mM MgCl_2_, 0.5 % Triton X-100, 30 % glycerol, and a protease inhibitor tablet (Roche)]. We lysed the cells for 2 min at 37°C and then collected the 300 µl lysis buffer containing the free tubulin fraction, and mixed with 100 µL 4x SDS sample buffer [200 mM Tris-HCl at pH 6.8, 8 % SDS, 40 % glycerol, 4 % β-mercaptoethanol, 50 mM EDTA and 0,08 % bromophenol blue]. The polymerised tubulin fraction present in the remaining material attached to the well was collected with 400 µL 1x SDS sample buffer. 40 µL of each fraction containing SDS sample buffer were prepared for Immunoblotting.

### Kinesore-dependent Microtubule binding assays for Kinesin-1

Endogenous kinesin-1 in cells is about 70 % inactive, cytosolic (Hollenbeck, 1989). In order to quantify Kinesore caused kinesin-1 binding to microtubules, the tubulin extraction assay as described above was performed. We incubated HeLa and Ptk2 cells with DMSO (0.1 or 0.2 %), or Kinesore (50 or 100 µM) for 15 min. After lysing the cells with extraction buffer, cytoplasmic inactive kinesin-1 was collected. The remaining fraction of kinesin-1 bound to microtubules were dissolved with 400 µL 1x SDS sample buffer. 40 µL of each fraction containing SDS sample buffer were prepared for Immunoblotting.

### SDS-PAGE and Western blot

Proteins were separated by SDS-PAGE (8 % gels) and then transferred to a nitrocellulose membrane with an iBLOT 2 Gel Transfer Device (ThermoFisher Scientific, IB21001). Nitrocellulose membranes were blocked for 1 h with 5 % dried milk resuspended in TBS-Tween 1 % and incubated over-night with primary antibodies: anti-alpha-tubulin (Sigma, T6074, 1:1000 dilution) or anti-UKHC (Santa Cruz Biotechnology, SC-133184, 1:1000 dilution). Membranes were washed three times with TBS-Tween 1 %, incubated for 1 hour with a secondary antibody anti-Mouse (GE Healthcare, NA9310, 1:5000 dilution), washed three times with TBS-Tween 1 % solution and revealed with an ECL Western blotting detection kit (Advansta) and with Fusion Solo Vilber Lourmat camera (Witec ag).

### Immunofluorescence microscopy

For localization of endogenous kinesin-1 after Kinesore treatment, HeLa cells were grown until 50-60 % confluency in 12 mm coverslips in 12-wells plates and treated with 0.1 % DMSO or 50 µM in Ringer’s buffer for 15 min at 37°C. Cells were rinsed once with warm PBS and cell extracted to remove cytosolic inactive kinesin-1 (see Kinesore-dependent Microtubule binding assays for Kinesin-1 section above) before fixation. HeLa and Ptk2 cells were fixed with 100% cold methanol for 4 min at -20°C, permeabilized and blocked with 0.1% Triton X-100 and 2% bovine serum albumin in phosphate buffered saline (PBS) for 30 min. Cells were incubated with primary antibodies [mouse anti-UKHC for kinesin-1 (Santa Cruz Biotechnology, SC-133184, 1:500 dilution), rabbit anti-alpha-Tubulin (Abcam, ab18251,1:1000 dilution) and rabbit anti-myc (Sigma-Aldrich, 06-549, 1:100 dilution)] and secondary antibodies [mouse and rabbit Alexa Fluor 488, 568 and 647] in blocking buffer in an humidified chamber for 1 hour. After each labelling step cells were washed three times with PBS. Finally, coverslips were dried and mounted using ProLong™ Diamont Antifade Mountant.

### Expansion microscopy

After fixation (see section above) Ptk2 cells were expanded isotropically 2x following Gambarotto et al., 2020 (Gambarotto et al., 2020) detailed protocol to visualize single microtubule polymers after 100 µM Kinesore treatment using rabbit anti-α-Tubulin (abcam; ab18251; 1:500 dilution).

### Statistical analysis

Statistical analysis was determined by two-tailed unpaired Student’s t test, or one- or two-way analysis of variance (ANOVA) using GraphPad Prism software v.8. P values less than 0.05 were considered statistically significant. Asterisks indicate the degree of significance, with *P < 0.05, **P < 0.01, ***P < 0.001, or ****P < 0.0001; ns, not significant.

### Simulations

In the simulations (see Figure S4, and Videos S2 and S3), a microtubule was represented by a dynamic linear lattice, where the tubulin dimers were represented as sites. Two states were possible for the sites, corresponding respectively to GTP- and GDP-tubulin dimers. Between these states transitions were possible: a GTP-site switched into a GDP-site with a rate k_Hyd_ and a GDP-site turned into a GTP-site at rate k_Rep_. The first of these transitions represented spontaneous hydrolysis of a GTP-site, whereas the second transition captured lattice repair by a GTP-tubulin dimer incorporation after the spontaneous loss of a GDP-tubulin dimer from the microtubule lattice. For this process, an effective description was used instead of a molecular description, because in our experiments, tubulin exchange was not resolved on molecular time and length scales. The rate of GTP-tubulin hydrolysis for an exchanged tubulin k_HydX_ can in general be different from the hydrolysis rate for GTP-tubulin that got incorporated into the microtubule by attaching the plus end.

The lattice as a whole was defined by two dynamic states: the growing state and the shrinking state. In the growing state, sites in the GTP state were added at rate k_A_ at one end, henceforth called the plus end. In contrast, the other end, called the minus end, was considered dynamically inert. This property was represented by the usage of stabilized seeds in the *in vitro* experiments. In the shrinking state, sites were removed one by one from the plus end at rate k_D_. The transition of the lattice from the growing to the shrinking state was correlated to a microtubule catastrophe with a rate k_Cat_. Because of the experimental inaccessibility of the tubulin dimer state, catastrophe was not restricted to occur when the final site was in the GDP state. Rescue events, corresponding to the transition from a shrinking to a growing state, were captured with a probability p_Res_ after the removal of a site if the new site at the plus end was a previously repaired site, that is, it was in a GTP state that resulted from a transition from a GDP state.

Molecular motors were described as particles hopping directionally on the lattice with a rate k_Hop_. These particles were able to move to the neighbouring site in the plus end direction if this neighbouring site was empty. This exclusion property accounted for steric interactions between molecular motors. It was assumed that a reservoir was formed by detached motors such that particles were attached to empty sites on the lattice at a constant rate k_MotorA_. Changes in this rate corresponded to changes of the motor concentration in the experiments. Particles were detached from the lattice at rate k_MotorD_.

The motor-induced exchange of GTP-tubulin for GDP-tubulin was captured by a probability p_X_ that a site in the GDP state was replaced by a GTP site as a particle hopped away from this site. Similar to the spontaneous exchange of a GTP-tubulin dimer for a GDP-tubulin dimer, this was considered to be an effective description of the potentially much more involved molecular process underlying the motor-induced exchange of a GTP-tubulin dimer for a GDP-tubulin dimer.

Simulations were carried out using a variant of the Gillespie algorithm, where initially the total rate k_tot_ of all possible processes was determined. After the time to the following event had been drawn from an exponential distribution with the characteristic time k_tot_ ^-1^, the identity of this event was determined according to the relative contribution of all possible events to the total rate. All the simulations were performed using a custom-made C program.

### Parameters

Whenever possible, the selected parameter values in the simulation were directly taken from our experiments (see Table S1). The parameters k_Hyd_, k_HydX_ and p_Res_ could not be inferred from our *in vitro* experiments, because the state of nucleotide bound to the tubulin dimer was inaccessible. The value of k_Hyd_ was taken from literature (Melki et al., 1996).The simulation results were insensitive to variations in k_HydX_ as long as this value was below a threshold value. The value of p_Res_ was used as a fit parameter to achieve in the simulations the rescue frequency observed in the *in vitro* experiments in absence of molecular motors and kept also for the simulations in presence of motor particles. The parameters associated with the dynamic of the particles representing the motors were all taken from experiments with the exception of p_X_. By keeping all the other parameter values constant, this value was fixed such that the same rescue frequency as in the 0.4 nM kinesin-1 experiment was produced by the simulation in steady state. For simulation corresponding to other motor concentration only k_MotorA_ value was changed in proportion to the motor concentration change. Explicitly, for a motor concentration c=λc_0_, where c_0_=1 nM, then k_MotorA_=λk_MotorA,0_, where k_MotorA,0_ is the motor attachment rate for the concentration c_0_ of kinesin-1.

**Table S1.** **Parameters used during the simulation with the values and the meaning corresponding to each parameter.** ^*^values measured in this work; ^#^value fitted to reproduce the rescue rate in absence of motors; ^†^value fitted to reproduce the rescue rate in presence of motors (0.4 nM kinesin-1).

| Parameter | Value | Significance |
| --- | --- | --- |
| $k_A$ | $120 \text{ min}^{-1}$ | attachment rate of tubulin at plus end<sup>*</sup><br>(corresponds to a polymerisation velocity of $0.96 \mu\text{m}/\text{min}$) |
| $k_D$ | $1620 \text{ min}^{-1}$ | detachment rate of tubulin at plus end<sup>*</sup><br>(corresponds to a de-polymerisation velocity of $12.96 \mu\text{m}/\text{min}$) |
| $k_{Cat}$ | $0.3 \text{ min}^{-1}$ | catastrophe rate<sup>*</sup> |
| $p_{Res}$ | 0.15 | probability of rescue if site at plus end is GTP<sup>#</sup> |
| $k_{Hyd}$ | $3 \text{ min}^{-1}$ | hydrolysis rate of tubulin incorporated at plus end (Melki et al., 1996) |
| $k_{Rep}$ | $6 \times 10^{-4} \text{ min}^{-1}$ | spontaneous tubulin exchange rate<sup>*</sup> |
| $k_{HydX}$ | $7.5 \times 10^{-3} \text{ min}^{-1}$ | hydrolysis rate of exchanged tubulin incorporated<sup>#</sup> |
| $k_{MotorA,0}$ | $1.98 \times 10^{-3} \text{ min}^{-1}$ | motor attachment rate to an empty tubulin dimer for 1 nM kinesin-1<sup>*</sup> |
| $k_{MotorD}$ | $5.4 \text{ min}^{-1}$ | motor detachment rate<sup>*</sup> |
| $k_{Hop}$ | $960 \text{ min}^{-1}$ | motor stepping rate<sup>*</sup><br>(corresponds to a motor velocity of $7.68 \mu\text{m}/\text{min}$) |
| $p_X$ | $1.8 \times 10^{-9}$ | probability to induce repair upon a motor step<sup>†</sup> |

## Supplementary Information

**Figure S1.**
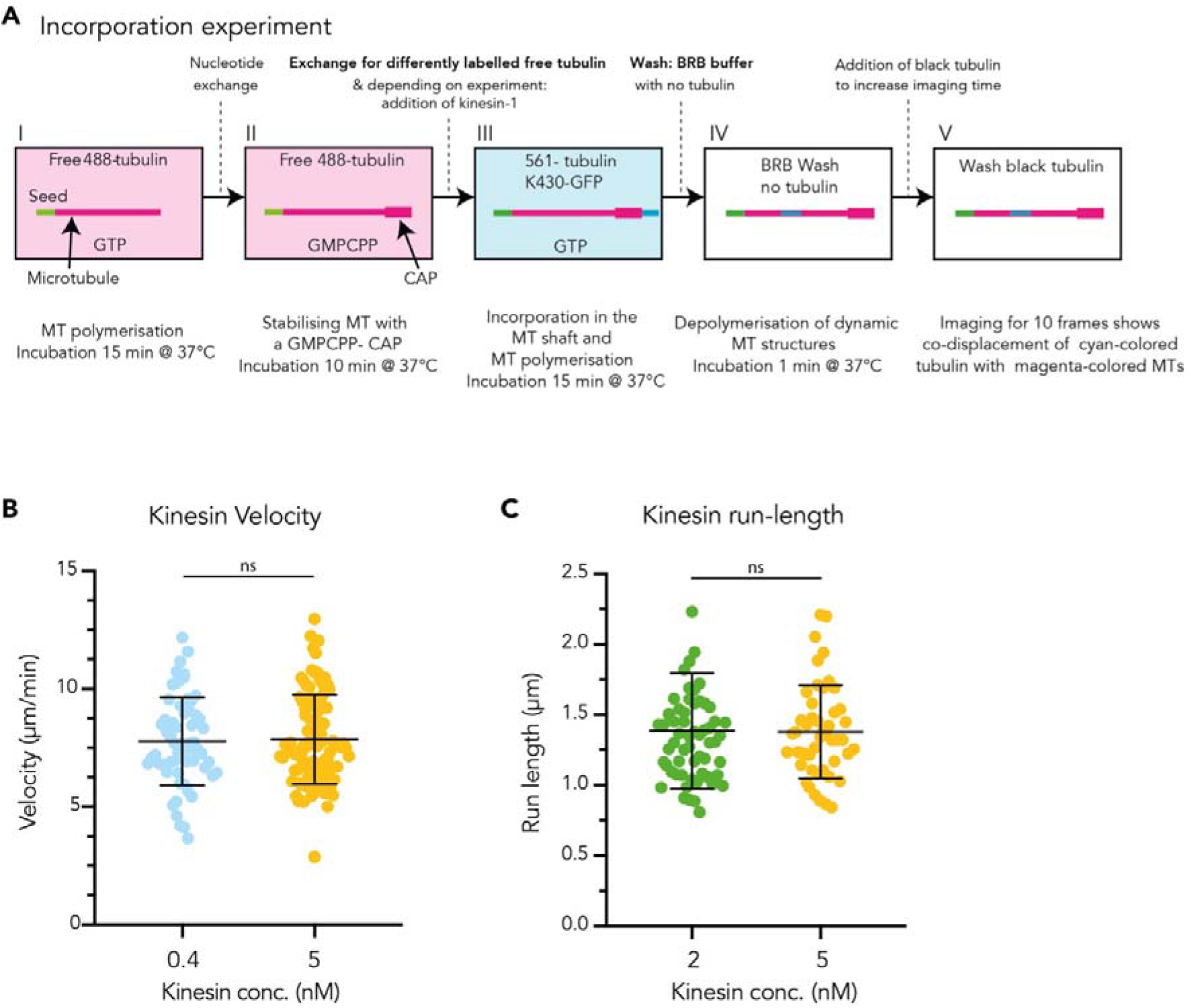
Incorporation assay and kinesin-1 dynamics. **(A)** Experimental setup of the incorporation assay. Flow chamber enables sequential exchange of solutions. **I** Microtubules polymerise form GMPCPP-seeds in the presence of tubulin 8 µM (20 % labelled), **II** Stabilizing GDP-microtubules with a GMPCPP-Cap. Exchange of nucleotide from GTP to GMPCPP and free tubulin 5 µM (70 % labelled), **III** Incorporation condition with 9 µM tubulin (100% labelled different colour) and kinesin-1 at different concentrations (0 nM, 1 nM, 2 nM, or 5 nM), **IV** Depolymerisation of all dynamic structures grown on top of the microtubule **in** BRB with no tubulin, **V** Stabilizing the microtubule for imaging with 8 µM black tubulin. **(B)** Kinesin-1 added to the incorporation assay moved along microtubules at an average velocity of 7.7 µm/min, comparable to published data (Furuta et al., 2013). Mean with SD of two independent experiments with a total of analysed kinesin-1 tracks: n=106 (0.4 nM), n=161 (5 nM); Statistics: unpaired T test; ns, p=0.7931. **(C)** Kinesin-1 run length in the presence of 8 µM tubulin at different kinesin-1 concentrations. Kinesin-1 run lengths were measured in 2 independent experiments for each condition with a total of n=163 (2 nM), n=69 (5 nM) tracks. Statistics: unpaired T test; ns, p= 0.9328. Mean with SD.

**Figure S2.**
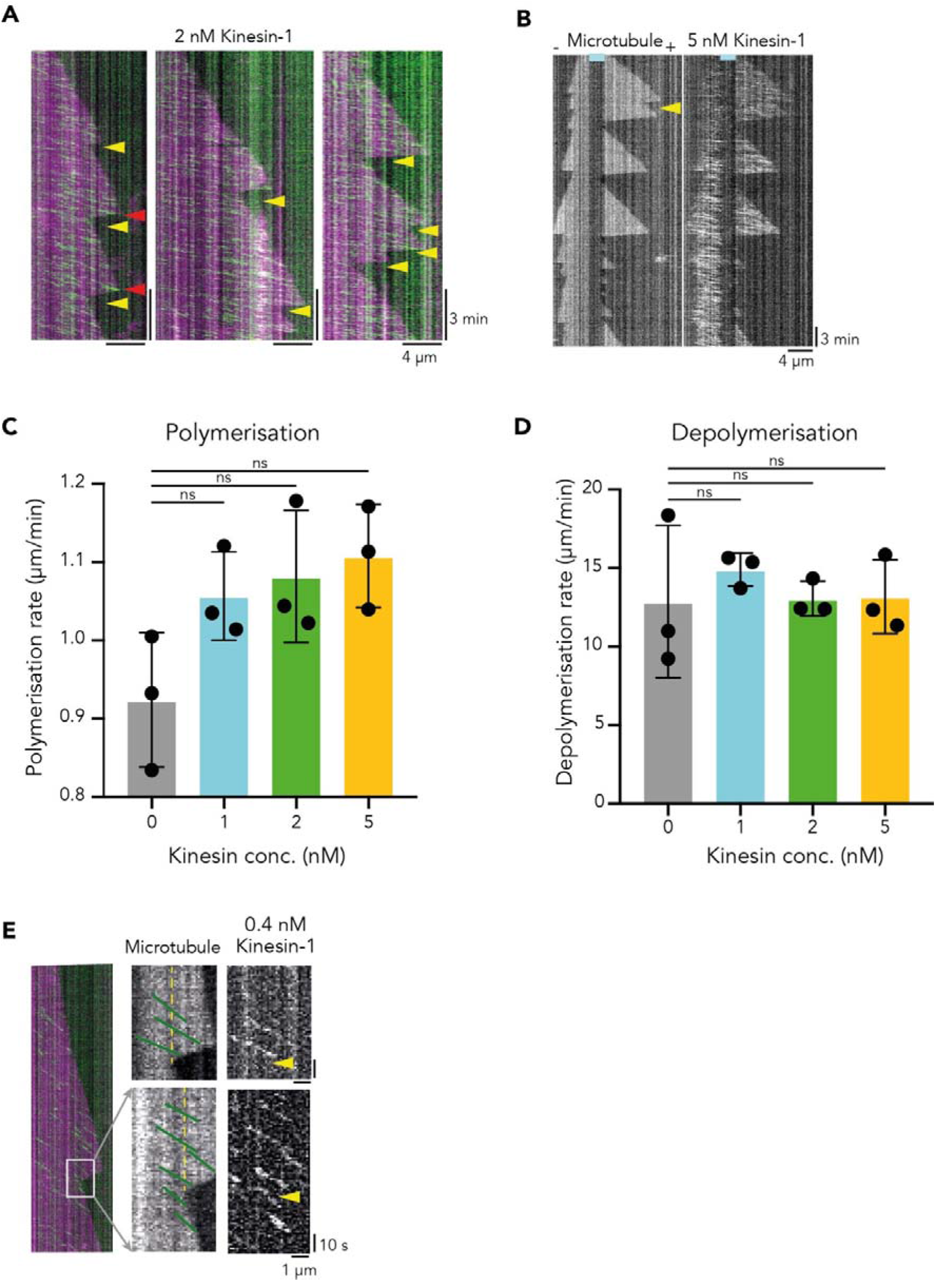
Microtubule dynamics in the presence of kinesin-1. **(A)** Representative kymographs of dynamic microtubules (8 µM tubulin, magenta) in presence of running kinesin-1 (2 nM, green). Yellow arrowheads indicate rescue sites. Red arrowheads indicate colliding of kinesin-1 which was not leading to rescue events or changing the microtubule depolymerisation behaviour. **(B)** Representative kymograph of a dynamic microtubule (8 µM tubulin) and corresponding kymograph of running kinesin-1 (5 nM). Yellow arrow indicates rescue site. Only few kinesins were found on the Taxol stabilized GMPCPP-seed (cyan line). Plus- and minus-ends are indicated. **(C)** Mean experimental polymerisation speed of microtubules in presence of different kinesin-1 concentrations of three independent experiments. Statistics: one-way ANOVA. Mean with SD. **(D)** Mean experimental depolymerisation speed of microtubules at different kinesin-1 concentrations. Statistics: one-way ANOVA. Mean with SD. **(E)** Representative microtubule kymograph (8 µM tubulin, magenta, left) in the presence of running kinesin-1 (0.4 nM; green). In microtubule kymograph (middle) kinesin-1 tracks are drawn as green lines, future rescue site is indicated by yellow dashed line. Single kinesins show no change in running behaviour at the future rescue site. Yellow arrowheads indicate rescue sites in kinesin kymograph (right).

**Figure S3.**
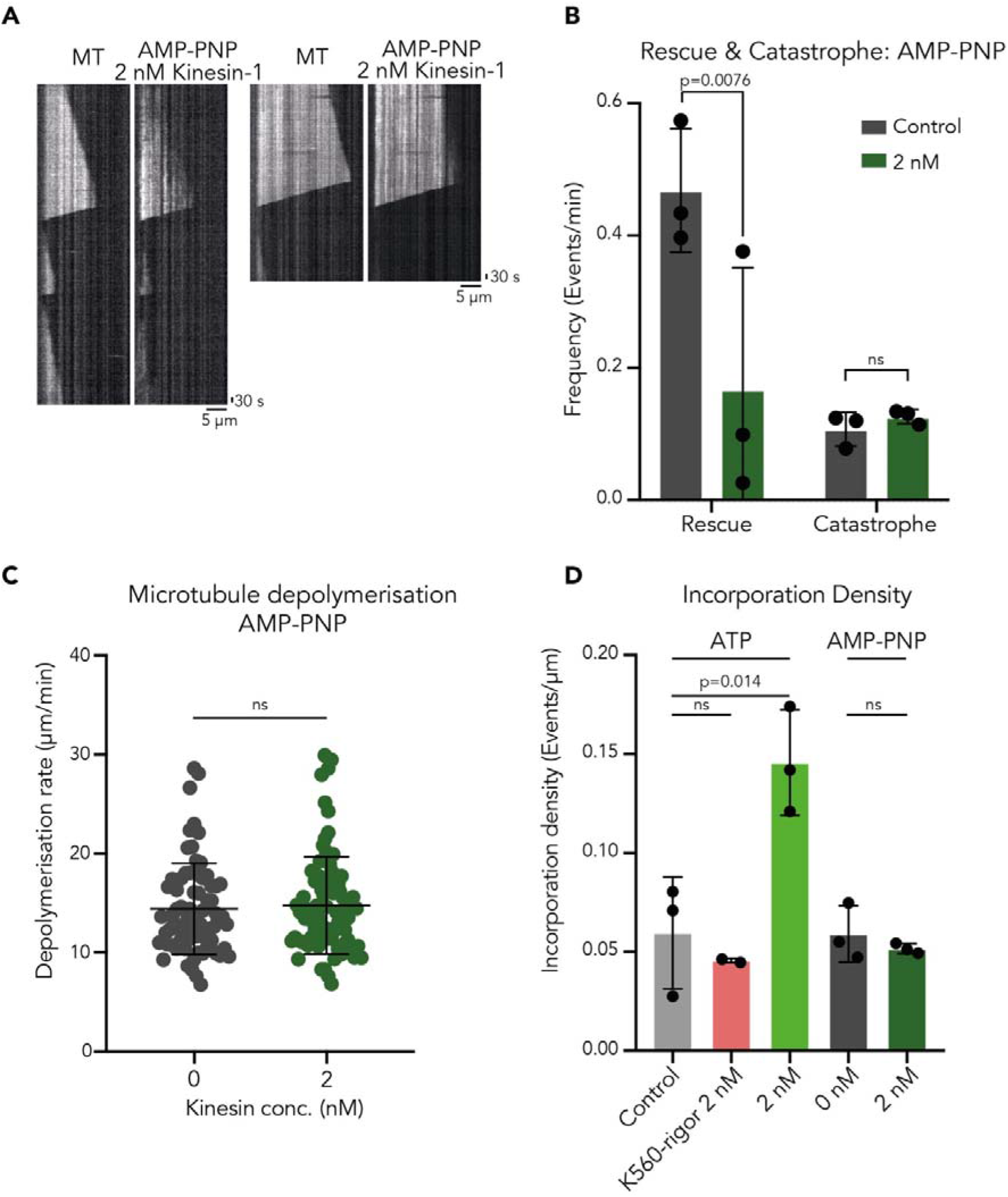
Only running kinesin-1 induces microtubule rescue and tubulin incorporation along the microtubule shaft. **(A)** Representative kymographs of dynamic microtubules (6.5 µM tubulin) and corresponding kymographs of immotile kinesin-1 (2 nM) in the presence of AMP-PNP. Immotile kinesin-1 densely covered the microtubule. **(B)** Rescue frequency in the absence of kinesin-1 (Control) and in presence of immotile 2 nM kinesin-1 (1 mM AMP-PNP). The rescue and catastrophe events were measured in three independent experiments for each condition, total number of microtubules analysed: n=152 (Control); n=162 (2 nM kinesin-1). Statistics: multiple t test. Mean with SD. **(C)** Microtubule depolymerisation did not change in the presence of immotile kinesin. Depolymerisation rates were measured in 3 independent experiments for each condition with a total of n=71 (0 nM), n=82 (2 nM) microtubules. Statistics: unpaired T test. Mean with SD. **(D)** Density of incorporation sites (total number of incorporation sites divided by total microtubule length) in control condition (ATP, 0 nM kinesin-1), 2 nM K560-rigor (ATP), 2 nM kinesin-1 (ATP), control (AMP-PNP, 0 nM kinesin-1), 2 nM kinesin-1 (AMP-PNP). Statistics: one-way ANOVA. Mean of experiment average with SD.

**Figure S4.**
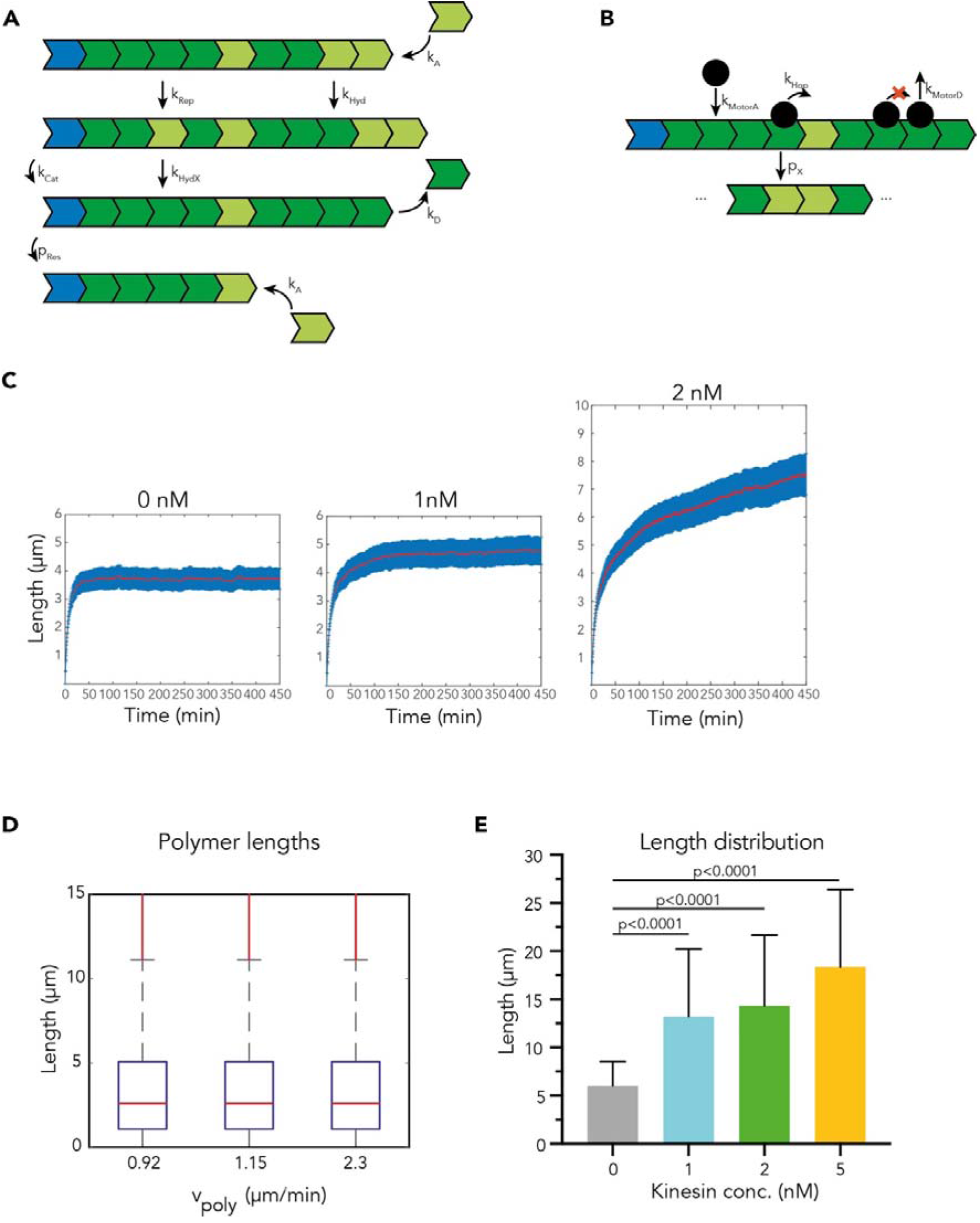
Kinesin-1 affects microtubule length. **(A)** Illustration of polymer dynamics indicating the different processes accounted for the simulation. GTP- or GDP-tubulin dimer were encoded as lattice sites (light green for GTP-tubulin dimer and dark green for GDP-tubulin dimer). The polymer seed is represented in blue. Dynamical parameters are highlighted (for values see Methods, Table S1). **(B)** Illustration of motor particles interacting with the polymer according to the different parameters encoded in the simulation. Motor particles are represented by black dots. Dynamical parameters are highlighted (for values see Methods, Table S1). **(C)** Numerical Simulations: Polymer length over time plotted for different motor particle concentrations: 0 nM (Control), 0.4 nM, 1 nM and 2 nM motor particles. Mean length (red) with SE (blue). The simulation time is longer compared to Figure 4A to show the asymptopic behaviour. **(D)** Numerical Simulations: Polymer length at different polymerisation speeds observed experimentally (Figure 2B) in presence of different kinesin-1 concentrations (0.92 µm/min for 0 nM, and 1.15 µm/min for 5 nM). A scenario with double velocity (2.3 µm/min) was also considered. Note that the microtubule length is not affected by the polymerisation speed. Median (red line) with 25^th^ and 75^th^ percentiles (blue box) are indicated. **(E) Mean** microtubule length observed experimentally in presence of different kinesin-1 concentrations at timepoint 30 min of three independent experiments. Statistics: one-way ANOVA. Mean with SD.

**Figure S5.**
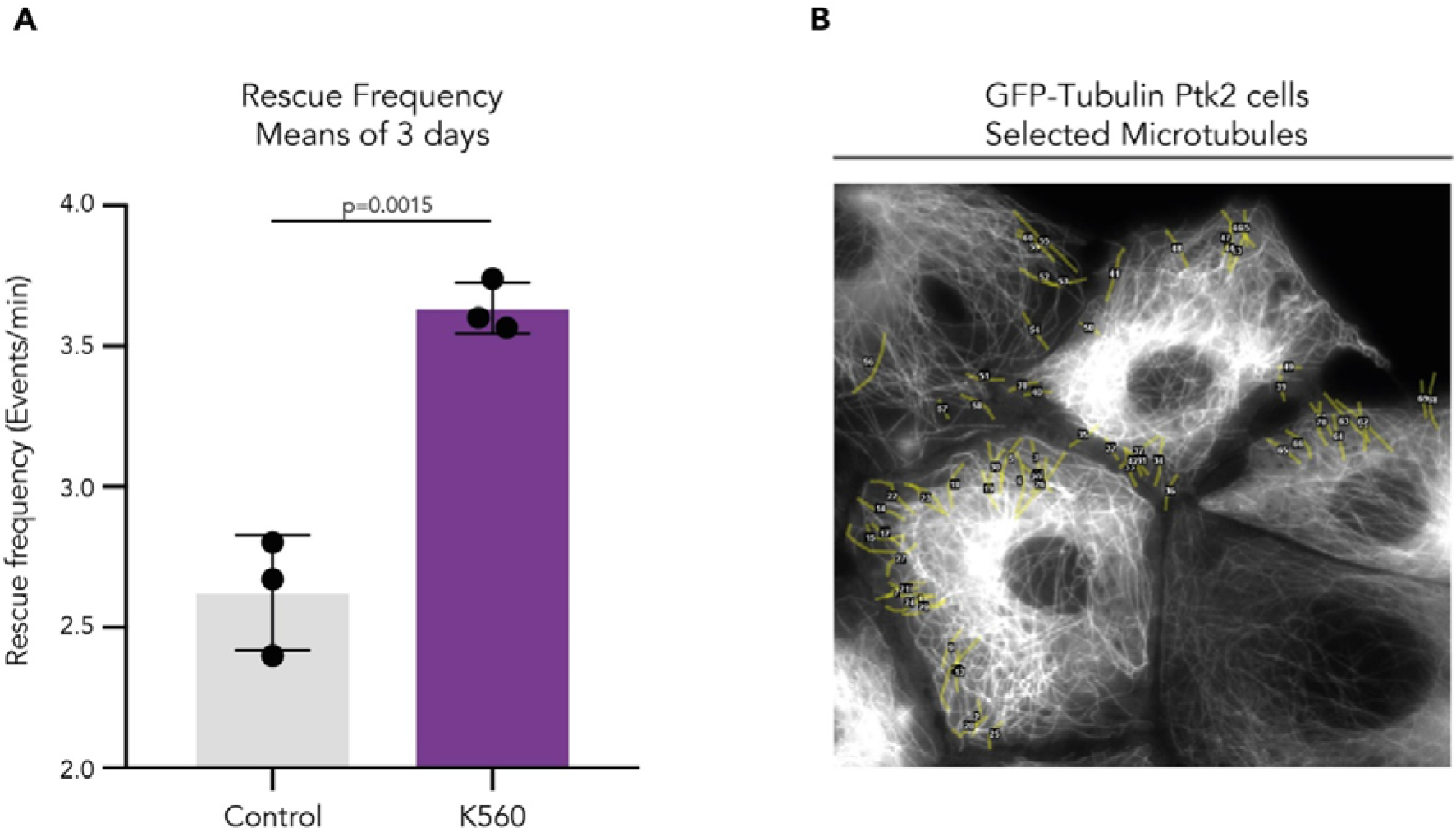
Kinesin-1 overexpression increases microtubule rescue frequency in cells. **(A)** Mean rescue frequency of GFP-Tubulin HeLa cells overexpressing K560-mCherry from three independent experiments. Statistics: unpaired T test. Mean with SD. **(B)** Representative image of selected microtubules in GFP-Tubulin Ptk2 cells for analysis. Scale bar: 10 µm.

**Figure S6.**
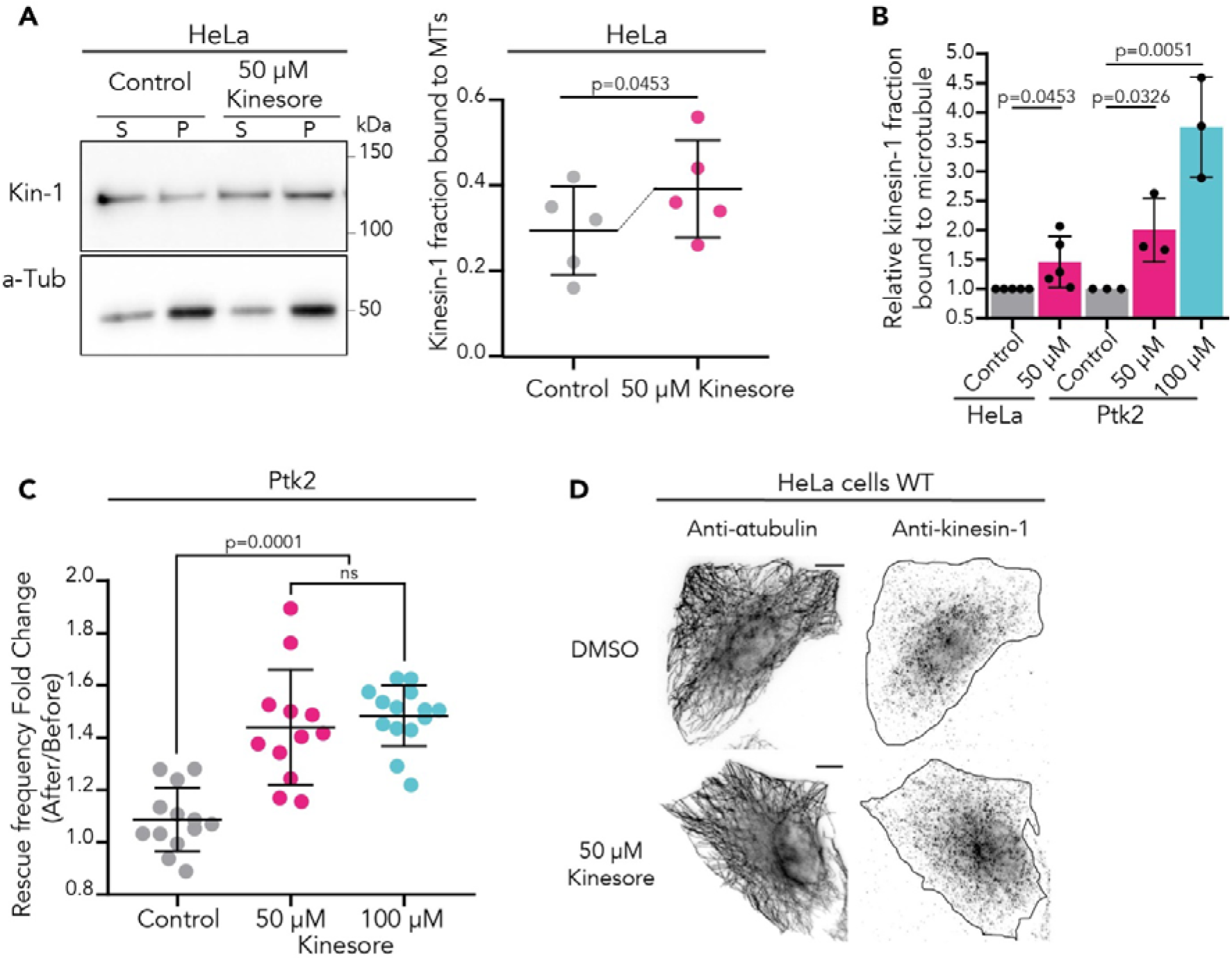
Increased fraction of kinesin-1 running on microtubule after Kinesore treatment. **(A)** Kinesore addition for 15 min increases kinesin-1 binding to microtubules in HeLa cells (MT, Pellet fraction, P). Representative Western Blot with quantification. Fraction of active kinesin-1 bound to microtubules determined by the Kinesore-dependent microtubule binding assays for kinesin-1 (see methods). Control (0.1 % DMSO) and 50 µM Kinesore treated HeLa cells. Statistics: unpaired T test. Mean with SD from four independent experiments. **(B)** Kinesore addition for 15 min increases kinesin-1 binding to microtubules. Relative binding of active kinesin-1 (bound to microtubules) after treatment with 0.1% or 0.2% DMSO (Control), 50 µM Kinesore, and 100 µM Kinesore in HeLa and Ptk2 cells. Statistics: unpaired T test. Mean with SD from three to four independent experiments. **(C)** Fold change of microtubule rescue frequency before and after Kinesore treatment in GFP-Tubulin Ptk2 cells. Cells were treated with 0.2% DMSO (Control; n=130 MT from 13 cells), 50 µM (n=120 MT, from 13 cell) and 100 µM Kinesore (n=140 MT from 14 cell) and the dynamics of single microtubules were analysed in three independent experiments. Statistics: two-tailed T test. Mean with SD. (D) Immunostaining of native kinesin-1 shows that Kinesore increases the binding of kinesin-1 to microtubules. Representative image of fixed HeLa cells treated with DMSO (Control) or 50 µM Kinesore for 15 min. Anti-α-tubulin and anti-kinesin-1 were used to visualise microtubule network and kinesin-1 distribution, respectively. Scale bar: 10 µm.

**Figure S7.**
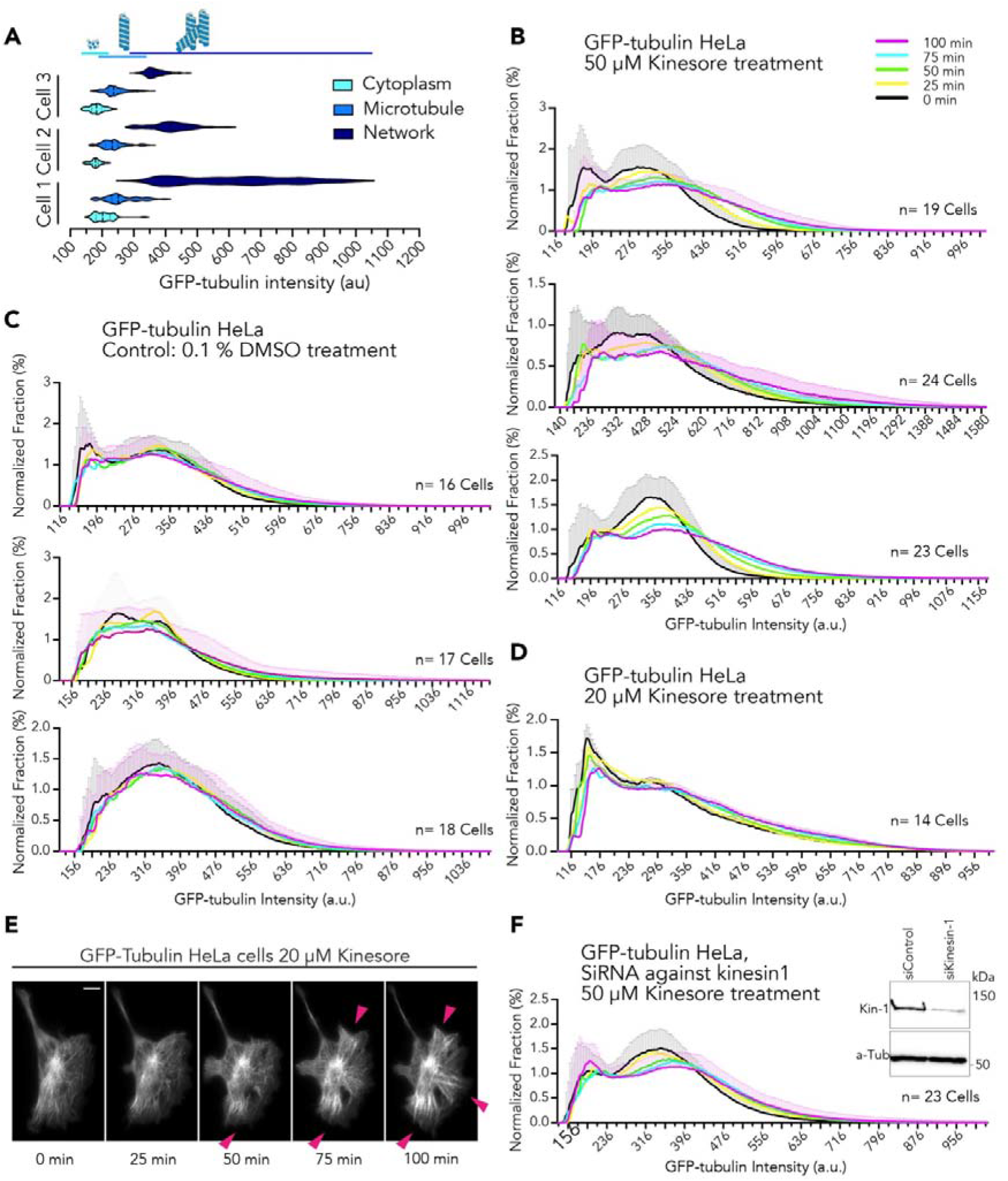
Kinesin-1 activity increases microtubule network and reduces free dimer pool. **(A)** Subdivision of the GFP-Tubulin fluorescence intensity in HeLa cells. Fluorescence intensity was measured at cytoplasm (tubulin dimers), along single microtubules, and in areas of denser microtubule network. Illustration above the graph shows the subdivision of fluorescence intensity into the three groups: tubulin dimer, single microtubules, microtubule network. Three cells from three independent experiments. Total measured grey values: cytoplasmic tubulin dimers (n=1504), single microtubule (n=2008), microtubule network (n=1608). Normalized fraction of the global GFP-Tubulin fluorescence intensity in HeLa cells. Cells were imaged every 5 min for 100 min after the addition of 0.1% DMSO to GFP-Tubulin HeLa cells (Control), or 20 µM, 50 µM Kinesore. The increment of the microtubule network is kinesin-1 dependent. DMSO does not affect the microtubule mass and the histogram curve is not shifting higher fluorescence intensities. **(B)** Kinesin-1 activation through 50 µM Kinesore shifts the histogram to higher fluorescence intensities over time. The microtubule mass is increasing and the free dimer pool decreasing. Three independent experiments shown with a total of 66 analysed cells. Normalized Mean with SD. **(C)** DMSO is not affecting the microtubule mass and the histogram curve is not shifting higher fluorescence intensities over time. Three independent experiments (here and in Figure 5B) shown with a total of 51 analysed cells. The legend is shown on the right. Normalized Mean with SD. **(D)** Kinesin-1 activation through 20 µM Kinesore shifts the histogram to higher fluorescence intensities over time, but less than 50 µM Kinesore. Total of 14 analysed cells. Normalized Mean with SEM. **(E)** GFP-Tubulin HeLa cells imaged every 5 min for 100 min after 20 µM Kinesore addition. Addition of Kinesore leads to reorganization of the microtubule network towards the cell rear where the network density increases (magenta arrowheads). **(F)** Kinesin-1 knockdown GFP-Tubulin HeLa cells after addition of 50 µM Kinesore. Corresponding Western blot of kinesin-1 expression levels in GFP-Tubulin HeLa cells knockdown for kinesin-1 (siRNA kinesin-1). Total of 23 analysed cells. Normalized Mean with SD.

**Figure S8.**
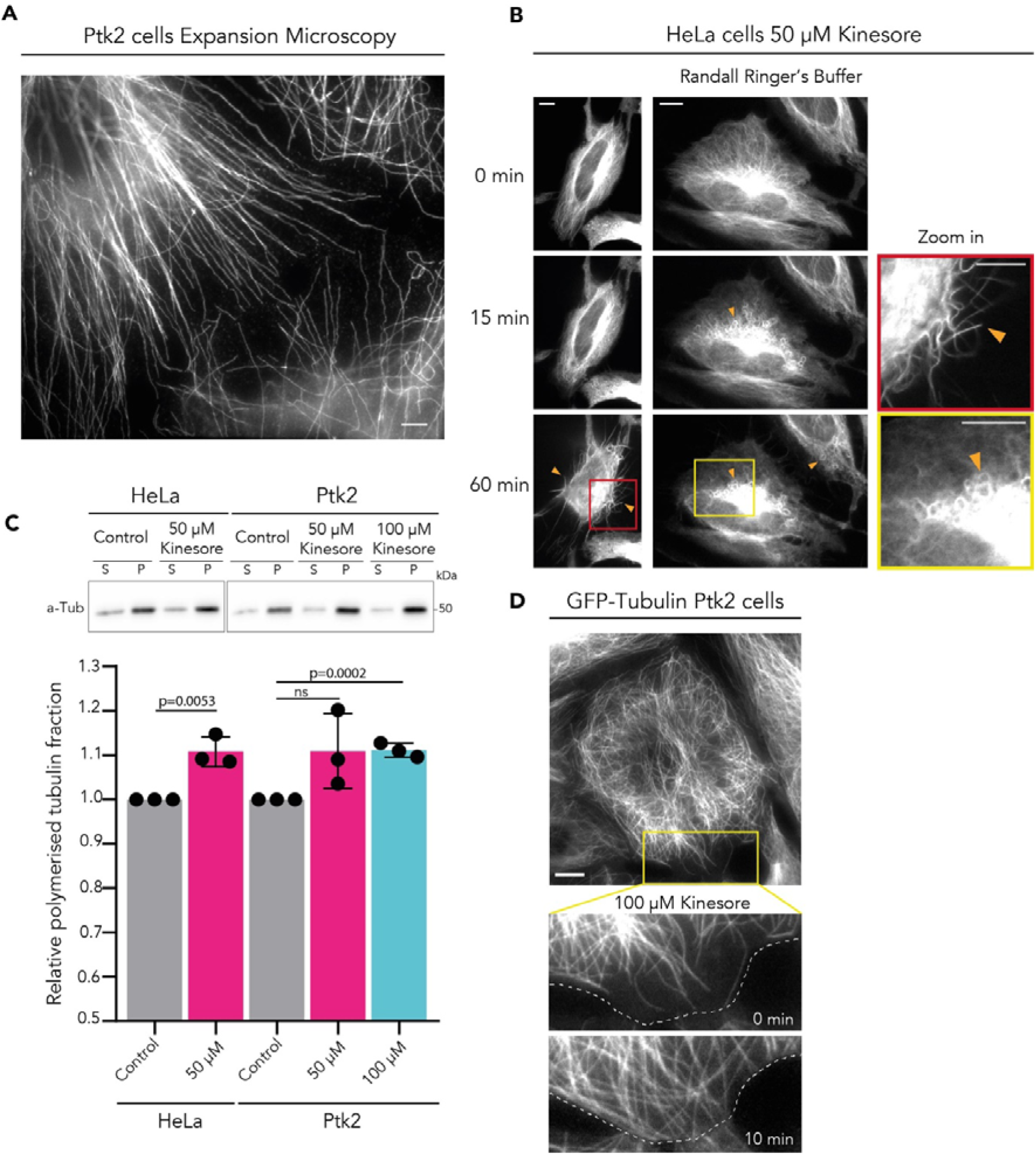
Kinesore affecting microtubule network organisation. **(A)** Ultrastructure expansion microscopy image of Ptk2 cells after 100 µM Kinesore treatment. A rabbit anti-α-tubulin was used to visualize the expanded microtubule network. Scale bar: 10 µm. **(B)** Representative images of GFP-Tubulin HeLa cells treated with 50 µM Kinesore during 0, 15 and 60 minutes in Randall Ringer’s Buffer. Yellow arrowheads and zoom-in red and yellow squares show microtubule bundles and loops reported previously in Randall et al., 2017. Scale bar: 10 µm. **(C)** Effect of Kinesore on microtubule mass. Representative Western Blot with quantification. HeLa cells were treated with 0.1% DMSO or 50 µM Kinesore, Ptk2 cells were treated with 0.2 % DMSO, 50 µM, and 100 µM Kinesore for 40 min at 37°C. Cells were lysed and microtubules (pellet fraction, P) were separated from the cytosolic tubulin dimer pool (supernatant fraction, S). The two fractions were analysed by Western blot. Statistics: unpaired T test. Mean relative tubulin levels present in the pellet with SD from three independent experiments. **(D)** The zoom-in (yellow square) shows a representative microtubule network reorganization after 100 µM Kinesore treated GFP-Tubulin Ptk2 cells (Video S7). The cell outlines, indicated with white dashed lines, help to show that the microtubule tips are closer to the plasma membrane after 10 min of Kinesore addition.

**Figure S9.**
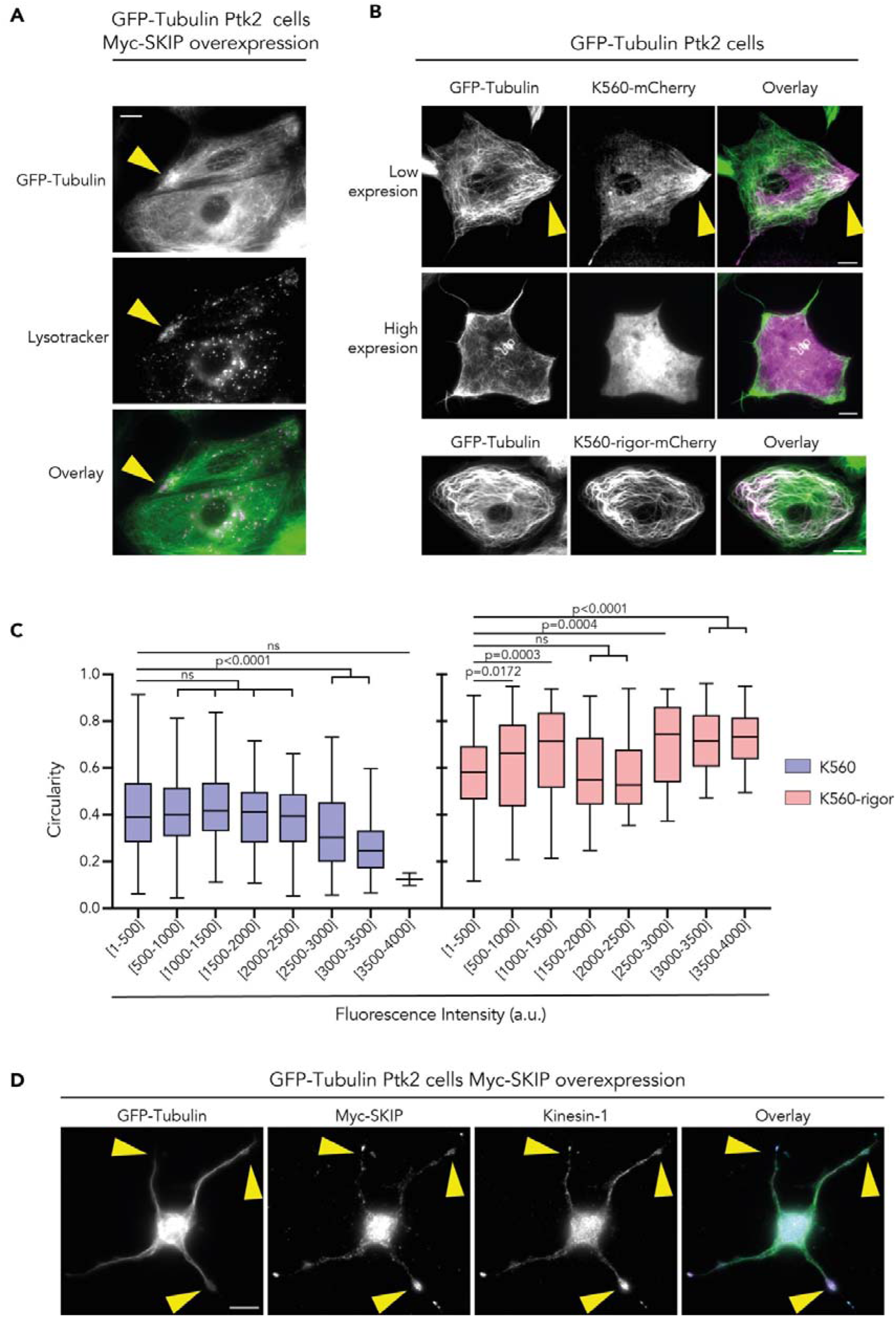
Kinesin-1 breaking cell polarity. **(A)** Representative TIRF image of GFP-Tubulin Ptk2 cell overexpressing myc-SKIP and stained with lysotracker to visualise lysosome movement. The yellow arrowhead highlights a region of microtubule network enrichment colocalizing with active lysosome transport. Scale bar: 10 µm. **(B)** Representative TIRF images of GFP-Tubulin Ptk2 cells transfected with a constitutively active mutant of kinesin-1, K560 (top). In K560 low expression cells, local increase of microtubule density was observed. The yellow arrowhead indicates a region of microtubule network enrichment colocalizing with kinesin-1. In K560 high expression cells, the MT architecture was completely altered. The ATPase rigor mutant of kinesin-1 (bottom) decorates and bundles microtubules since it binds microtubules but cannot walk along them. Scale bar 10 µm. **(C)** Cell circularity of GFP-Tubulin Ptk2 cells overexpressing K560 compared to GFP-Tubulin Ptk2 cells overexpressing K560-rigor. Cell circularity is plotted over binned mean fluorescence intensity. K560 (n=67 cells) and K560-rigor (n=58 cells) from three independent experiments. Statistics: one-way ANOVA. Boxplots show median with the 25th and 75th percentile as box edges. **(D)** Fixed GFP-Tubulin Ptk2 cells overexpressing myc-SKIP with kinesin-1 endogenous localization. Yellow arrowheads indicate regions of microtubule network enrichment colocalizing with kinesin-1 and lysosome transport. Scale bar: 10 µm.

**Video S1**.

Immotile kinesin-1 (2 nM, magenta) bound to dynamic microtubule (8 µM, green), does not change the microtubule depolymerisation behaviour.

**Video S2**.

Simulation of polymer dynamics in the absence of motor particles over 45 min. The seed is represented in blue. In the polymer the GDP-tubulin dimers are represented in dark green and the GTP-tubulin dimers in light green.

**Video S3**.

Simulations of polymer dynamics in the presence of 2 nM motor particles over 45 min. The seed is represented in blue. In the polymer the GDP-tubulin dimers are represented in dark green and the GTP-tubulin dimers in light green. Motor particles running along the polymer are represented in black.

**Video S4**.

Microtubule (8 µM tubulin) length in the absence of kinesin-1 over 45 min imaged every minute.

**Video S5**.

Microtubule (8 µM tubulin) length in the presence of 5 nM kinesin-1 over 45 min imaged every minute.

**Video S6**.

Microtubule network organization before and after 100 µM Kinesore incubation in GFP-Tubulin Ptk2 cells. Zoom-in of microtubules close to the plasma membrane. Distance between microtubule tips and plasma membrane is reducing after 10 min of Kinesore addition. The effect is enhanced over time.

**Video S7**.

Microtubule network organization before and after 100 µM Kinesore incubation in GFP-Tubulin Ptk2 cells. Zoom-in of microtubules close to the plasma membrane. Distance between microtubule tips and plasma membrane is reducing after 10 min of Kinesore addition. The effect is enhanced over time.

**Video S8**.

Heterogenous spatial K560-mCherry distribution impacts on cell morphology. Imaging of GFP-Tubulin Ptk2 cells after 8 hours of K560-mCherry overexpression for 40 hours. In order to visualize K560-mCherry distribution over time, the brightness in the m-Cherry channel was adjusted between frames of the movie (adjustment is indicated with yellow arrowhead).

